# Mix-it-up: Accessible time-resolved cryo-EM on the millisecond timescale

**DOI:** 10.1101/2025.07.22.666177

**Authors:** Lauren Alexandrescu, William Lessin, Gabriel C. Lander

## Abstract

Biological reactions often involve macromolecules that undergo substrate-induced conformational changes in under a second, yet capturing these transient states remains challenging. While high-resolution structural techniques such as X-ray crystallography and cryo-electron microscopy (cryo-EM) have advanced our mechanistic understanding of protein-substrate interactions, traditional sample preparation methods are too slow to capture rapid biochemical events. Time-resolved cryo-EM has emerged as a promising approach to visualize structural dynamics on microsecond-to-millisecond timescales, but its widespread adoption has been limited by costly equipment and challenges in achieving rapid mixing, application, and vitrification of samples in a reproducible manner. To address these limitations, we developed “Mix-it-up” (MIU), a modified spray device designed for rapid on-grid mixing and vitrification of cryo-EM samples. By manually applying one sample onto the EM grid, blotting, and subsequently spraying the second sample, we achieve on-grid mixing with a vitrification delay of as low as ~120 milliseconds. We demonstrate MIU’s time-resolved capabilities through high-resolution structure determination of mixed samples, pH-induced viral capsid contraction, and ligand-dependent complex formation. These findings establish MIU as a cost-effective, versatile tool for studying rapid biochemical processes and lay the groundwork for future applications to time-resolved cryo-EM.

## Introduction

Biological processes within and outside of cells involve dynamic, multi-component pathways, wherein macromolecules interact with cofactors and substrates while undergoing rapid conformational changes. Structural biology techniques such as crystallography and single-particle cryo-electron microscopy (cryo-EM) have provided high-resolution structural insights into macromolecular complexes, enabling detailed investigation of molecular interactions and the mechanisms of biochemical reactions^1–5^. While crystallographic structures are conformationally constrained by the requirements of crystal packing^6^, cryo-EM offers the capacity to visualize the conformational landscapes of macromolecules through rapid freezing and vitrification of aqueous samples^7,8^. However, conventional blot-based methods commonly used for single particle cryo-EM sample preparation are carried out on timescales of seconds to minutes^9,10^. These relatively prolonged timescales are more conducive to observing low-energy post-equilibrium states of biochemical reactions than the intermediate states associated with molecular recognition and binding or initial states of catalysis and processing. Reducing the timescales required to mix biological samples, substates, and reagents can enable visualization of the initial interactions and molecular events that trigger important biological mechanisms^11–15^.

Time-resolved cryo-EM has emerged as an attractive area of development for the cryo-EM community, as it could enable visualization of key time-dependent states associated with biochemical reactions. In order to achieve this feat, how-Å ever, the sample preparation steps must be performed on the timescale of these reactions, which typically occur on the order of microseconds to milliseconds^12,16–18^. Further, the enzymes and substrates involved must be thoroughly mixed prior to vitrification^19–21^. Several devices are currently under development to perform rapid sample application using spray or inkjet technologies, but these devices require high-cost equipment and do not incorporate a method for achieving sample mixing, thus preventing their use in time-resolved cryo-EM studies^16^. To date, time-resolved cryo-EM studies have shown limited application for biological samples due to lack of widespread equipment availability, technology usability, or reproducibility.

Previously, a low-cost, high-speed, ultrasonic spray-based vitrification device, “Shake-it-off”^22^, which was superseded by “Back it Up” (BIU)^23^, demonstrated its ability to prepare samples for high-resolution cryo-EM structure determination on a timescale of tens of milliseconds. Compared to commercially available devices for high-speed sample preparation that can cost over $1 million and are unable to apply multiple samples to a grid, the BIU costs less than $1000 to build, and the design of the custom parts have been made open source. The BIU device incorporates a piezoelectric transducer (hereafter referred to simply as a “piezo”) that enables aerosolization of a sample droplet when a high voltage is applied^22,23^. BIU is controlled by a Raspberry Pi computer running a customized, open-source Python program that automates the spraying and plunging steps. This device was shown to be capable of high-resolution structure determination, evidenced by a ~2 resolution map of apoferritin^23^. However, incorporation of mixed samples was not investigated in the study. Our work builds on this framework, implementing a “blot-then-spray” methodology^24,25^ for multi-sample mixing. Using this updated method, one sample is applied manually to the grid and the blotted automatically for a specified period of time, after which a secondary sample is ultrasonically dispensed to induce on-grid mixing (**Fig 1, Fig S1**). The time from application of the secondary sample to vitrification is ~120-720 ms. As a proof of principle, we determine high-resolution structures of apoferritin and aldolase from a mixed sample, capture pH-induced conformational changes of CCMV, and demonstrate ATP-induced complex formation of GroEL/ES. We term the third iteration of this ultrasonic spray technology “Mix-it-up” (MIU), following the basic design of the previous Shake-it-off and BIU implementations^22,23^.

**Figure 1.**
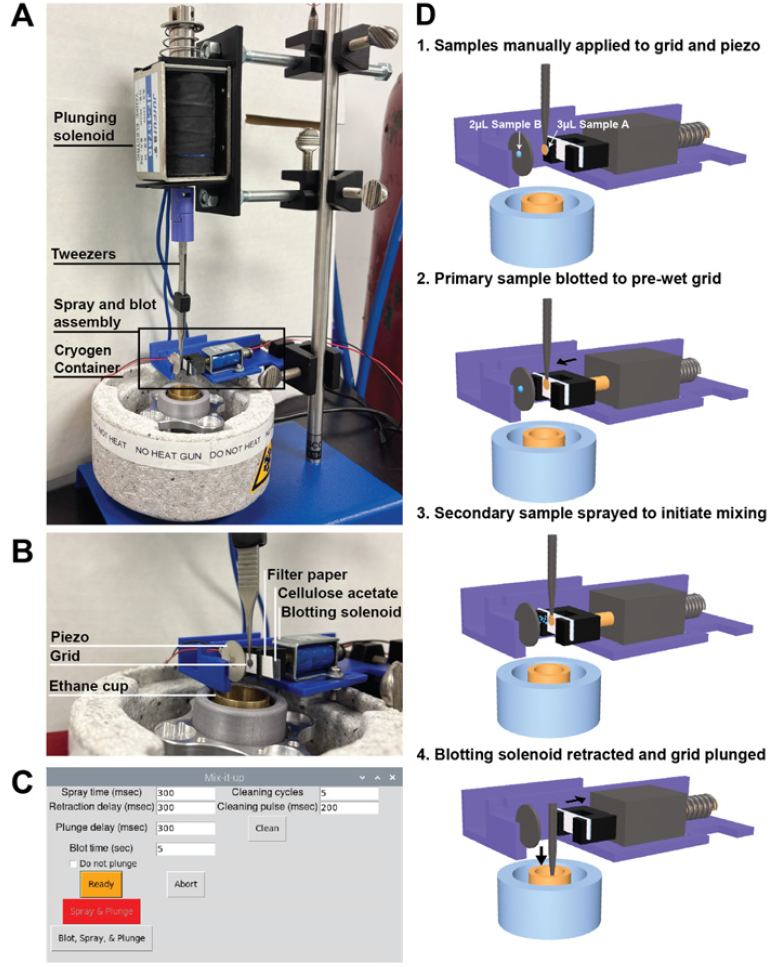
Mix-it-up design and method. **A**) The “Mix-it-Up” (MIU) assembly is based on the previously described “Back-it-up” (BIU) design^23^. The grid and tweezers are connected to a plunging solenoid positioned in front of the spray and blot assembly, positioned above the cryogen container. **B)** The spray and blot assembly consists of a piezo electric transducer (piezo) that aerosolizes an applied droplet of sample, directing a spray towards a grid positioned in front of a strip of filter paper used for blotting. **C)** The MIU GUI includes updates to the BIU GUI, including a parameter for blotting time and a button that triggers the blot-spray-and-plunge sequence. **D)** The method used for high-speed sample mixing includes the following steps: 1) manual application of a primary sample to the grid and a secondary sample to the piezo, 2) blotting of the primary sample to pre-wet the grid for improved mixed sample distribution, 3) initiation of spray to apply the secondary sample and trigger on-grid mixing, and 4) retraction of the blotting solenoid and plunging into the cryogen bath.

## Results

### Single-sample blotting with MIU is effective for high-resolution structure determination

BIU was designed for single-sample cryo-EM experiments, where sample is manually applied to a piezo, followed by automated aerosolization and application to the grid^23^. In the original BIU framework, through-grid wicking is used instead of conventional blotting to rapidly thin the sprayed sample prior to vitrification. After its publication, the BIU architecture was successfully adapted for conventional blotting for use in a light-coupled, time-resolved cryo-EM platform^26^, and in time-resolved studies of kainite receptors^27^. However, in our studies using the BIU design, we found through-grid wicking to be ineffective at reproducibly thinning the sprayed sample across EM grids, leading to regions of thick ice surrounded by limited regions of ice that were amenable to imaging, but nonetheless too thick to yield optimal particle distributions for routine high-resolution particle reconstruction (**Fig 2A**).

**Figure 2.**
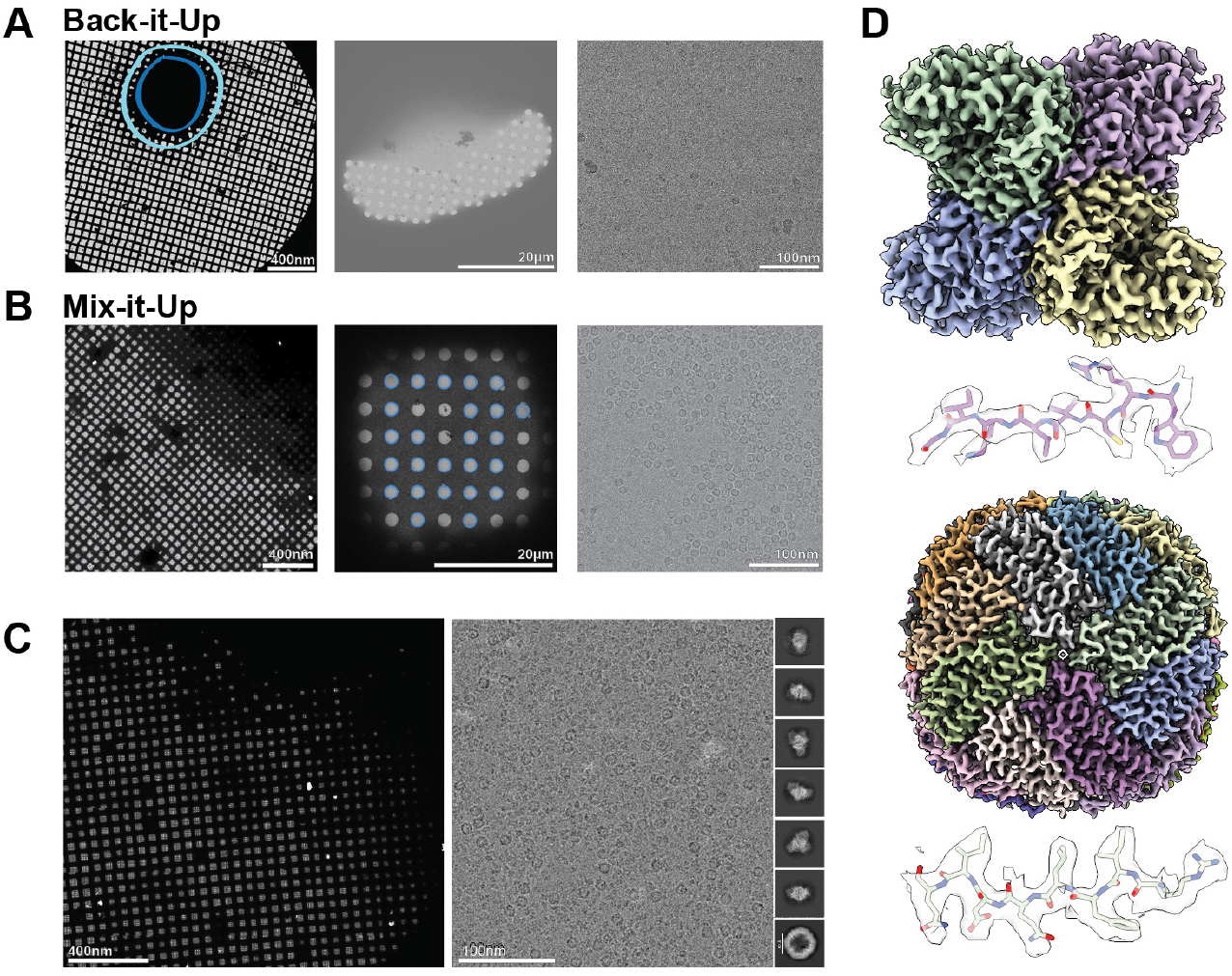
Improved ice distribution of mixed samples using MIU. **A, B)** Representative atlas (left), square (center), and high magnification images of apoferritin (right) obtained with BIU (A) and MIU (B), demonstrating the ice and particle distributions achieved with each method. A dark blue circle indicates regions of thick ice, and a light blue circle indicates regions of usable squares for targeting on the leftmost panel of (A). **C)** The atlas (left), representative micrograph (right), and representative 2D class averages obtained from the preparation of mixed apoferritin and aldolase samples using MIU. **D)** EM maps of aldolase (top) and apoferritin (bottom), both resolved to 2.9 Å, with representative peptide density shown beneath each map.

To adapt the framework for mixed-sample studies, we implemented a blot-then-spray methodology combining conventional blotting and spraying to promote on-grid mixing while achieving optimal ice thickness (**Fig 1, Fig S1**). We hypothesized that by using conventional blotting to generate a thinned layer of the primary sample across the grid, a secondary sample could be sprayed onto the wet surface, where it could rapidly spread and diffuse into the primary sample. We surmised that the small size of the ultrasonically dispensed droplets would not substantially increase ice thickness, and the thinness of the primary sample (tens of nanometers) would accommodate rapid diffusion of the secondary sample, which would in turn promote interaction between the samples.

To more effectively thin the sample for optimal ice thickness across the EM grid, we modified the blotting assembly to recapitulate the force distribution that would be introduced by manual blotting. In our studies, the circular design of the paper used in BIU resulted in inconsistent ice thickness across the grid due to an uneven distribution of force applied. To mitigate this effect, we designed a “clip” with two pillars on either end of a rectangular base that holds a strip of Whatman no. 1 filter paper securely in front of an equally sized strip of cellulose acetate (**Fig S1D**). This design aligns the point of contact of the filter paper and grid with the axis of movement of the blotting solenoid such that an even force is applied across the grid during the blotting process.

We first tested the capacity of the blot-then-spray method and updated framework to prepare cryo-EM grids that are amenable for single particle reconstructions. A homemade gold grid fenestrated with 2.2 µm holes spaced 2.2 µm apart was plasma cleaned and held by tweezers connected to the plunging solenoid. A volume of 3 µL of buffer was manually pipetted onto the side of the grid facing the blotting paper, and 2 µL of apoferritin was applied to the back of the piezo. Blotting was initiated with the “Ready” button on the graphical user interface (GUI), which triggers the forward movement of the blotting solenoid, positioning the filter paper in contact with the droplet of buffer. After three seconds, a 300 ms spray was triggered, immediately followed by simultaneous retraction of the blotting assembly and the grid was plunged into a liquid/solid ethane slurry. The time between the blotting paper retraction and the sample entering the ethane was measured to contribute an additional ~23 milliseconds.

The resulting atlas exhibited a gradient of ice thickness similar to that of grids prepared using a manual plunging device or automated blotting instruments such as the ThermoFisher Vitrobot (**Fig 2B**). Greater sample thinning was observed within the squares compared to that of preparations with the original BIU framework, yielding improved particle distribution of the sprayed apoferritin (**Fig 2B**). This proof-of-principle demonstrated the ability of our MIU device to generate cryoEM grids having a consistent distribution of thin, vitreous ice containing sprayed particles for high-resolution reconstruction.

### High-resolution structures of mixed apoferritin and aldolase samples using MIU

Previous studies demonstrated the effectiveness of “on-grid mixing,” a phenomenon in which samples are independently applied to a grid and mix upon contact within the foil holes^16,18,21^. We sought to demonstrate the ability of MIU to enable on-grid mixing by manually applying a primary sample to a grid, blotting, and subsequently spraying a secondary sample onto the grid (**Fig 1D**). Given that we observed sprayed apoferritin distributed across a cryo-EM grid that had been pre-wetted with buffer, we next aimed to characterize the sprayed apoferritin distribution across a grid that is pre-wetted with an aldolase sample.

To promote on-grid mixing, it is important to first consider the directionality of the grid during attachment to the plunging tweezers and sample application. The “front” of the grid, also referred to as the “sample side,” corresponds to the side where the holey foil has been applied to the grid, where it is relatively flat across the grid bars (**Fig S2A-B**). The “back” side of the grid contains deep wells between the grid bars and the foil (**Fig S2C**), creating an area where sample can pool, making it difficult to generate a thin layer of sample even when blotted from this side, and increasing the likelihood of creating regions of thick ice that are not suitable for imaging. To prevent this, the sample side of the grid faces the blotting assembly, as the manually applied primary sample has a larger bulk volume compared to the sprayed droplets of the secondary sample. The spray is therefore targeted at the back side of the grid, where the droplets are expected tospread into sample-filled holes of the fenestrated film rather than pool. Due to the dual-sided sample application in our blot-then-spray method, both sides of the grid are plasma cleaned to ensure hydrophilicity and promote adherence and mixing of the sprayed droplets with the layer of manually applied sample.

For our mixed apoferritin and aldolase studies, a 3 µL droplet of aldolase was first manually applied to the sample side of the grid, as described in the single-sample study. A 2 µL droplet of apoferritin was then applied to the back side of the piezo. Next, the blotting assembly was set to the forward position to initiate blotting of the aldolase sample. After five seconds of blotting, the spray was triggered, resulting in aerosolization of the apoferritin droplet. Apoferritin was sprayed onto the back side of the grid for 300 ms, during which the filter paper remained in contact with the grid. Following the spray period, the blotting assembly was retracted, and the grid was rapidly plunge frozen.

The resulting atlas demonstrates ice distribution was similar to that obtained in the single-sample experiment, with several pockets of thick ice as well as areas of thinner ice radiating outward from these clusters (**Fig 2C**). Notably, high magnification imaging of this grid revealed an uneven amount of mixing across the grid. The acquired micrographs could be categorized into several phenotypic groups, containing: 1) high aldolase particle concentration with no apoferritin particles present (**Fig S3A**), 2) high aldolase particle concentration with apoferritin particles present at low concentration (**Fig S3A**), 3) optimally mixed aldolase and apoferritin particles (**Fig S3B**), and 4) high apoferritin particle concentration with aldolase particles present at low concentration (**Fig S3C**).

The first case, in which only aldolase particles are observed in micrographs, is likely indicative of regions where aerosolized droplets of the sprayed sample did not contact the grid. Micrographs that exhibit a high concentration of aldolase but low apoferritin concentration could have resulted from 1) interaction of a small aerosolized droplet with the grid that did not spread across a large area, 2) the application of several individual droplets that did not pool following application to the grid leading to low, local concentrations of apoferritin, or 3) imaging at regions at the outer boundary of a sprayed droplet or pooled collection of droplets. Conversely, in micrographs that exhibit a high apoferritin concentration but low aldolase concentration, apoferritin appears to displace aldolase from the region in order to engage in preferred interparticle interactions, in some cases resulting in hexagonal particle packing. The micrographs with high apparent apoferritin concentrations may result from large droplets containing high amounts of apoferritin hitting the grid, or from many droplets hitting the grid with close proximity and pooling, leading to a localized increase in particle concentration.

While apoferritin and aldolase are readily distinguishable in our micrographs due to their large size and distinct structures, this ability to visually evaluate sample mixing will not extend to sprayed small molecules or buffer components that are often used to modulate biologically relevant reactions. Further, given the variation in the distribution and local concentrations of sprayed apoferritin throughout our dataset, we concluded that it was crucial to track the application of sprayed secondary samples that consist solely of buffer components, ligands, or small peptides, which cannot be resolved using single-particle cryo-EM. Thus, we incorporated 5nm gold fiducials into the sprayed sample of subsequent experiments in order to confirm areas of application on the grid. This approach has previously been used in foundational time-resolved EM studies to track the location of sprayed sample^28^. Nanogold fiducials are well-suited as high-contrast and readily identifiable markers that can be observed at high magnifications but may require computational deletion from the images for small or challenging particles due to artifacts that can occur during the analysis of images containing nanogold.

Processing of our mixed sample dataset yielded a ~2.9 Å resolution map of apoferritin and a 2.9 Å resolution map of aldolase (**Fig 2D, Fig S4, Table S1**). These data demonstrate the capacity of MIU to enable high-resolution structure determination of multiple samples mixed on-grid using our blot-then-spray methodology.

### pH-induced contraction of CCMV observed on the millisecond timescale

To assess whether MIU could capture conformational changes that take place on the millisecond timescale, we examined the well-established model system of the Cowpea chlorotic mottle virus (CCMV). CCMV is a plant virus of the Bromoviridae family that infects the cowpea species. The capsid has T=3 icosahedral symmetry, consisting of 180 identical copies of the coat protein^29^. During infection, CCMV capsids expand to release genetic material as they transition from the low-pH extracellular environment to higher pH conditions within the cellular cytosol. Both the contracted and expanded states of the capsid have been characterized by crystallography and cryo-EM^17,29^. At a pH of 4.8, or in the absence of divalent cations such as Ca^2+^ or Mg^2+^, the capsid favors a contracted form with a diameter of 29 nm^29^. At pH 7 or higher and under conditions of low ionic strength, the capsid expands to a diameter of 32 nm^17,29^. Notably, this shift in conformation is reversible with purified capsid, so that a sample can be expanded or contracted in vitro by changing the pH of its buffer.

A recent study implemented specialized hardware for rapid melting and re-vitrification of cryo-EM samples to study the timescale of CCMV capsid contraction following a photoacid-induced pH shift in the surroundings^17^. With a temporal resolution of 30 microseconds, the CCMV capsids were found to undergo a ~10 Å decrease in diameter, as well as a slight rearrangement in the capsid organization, corresponding to a partially contracted state.

In contrast, we focused on using MIU to trigger the full conformational change of CCMV capsids under low pH conditions to demonstrate MIU’s capabilities as a low-cost method for studying large-scale conformational changes on the millisecond timescale. To induce a pH shift and trigger the contraction of expanded CCMV capsids, we sprayed low pH buffer (pH 3.6) onto a grid pre-blotted with expanded CCMV capsids. Nanogold fiducials were mixed with the low pH buffer to track the localization of the buffer on the grid and confirm areas of sample mixing.

Our expanded CCMV control was prepared by dialyzing CCMV capsids stored in low pH buffer (pH 4.6) into high pH buffer (pH 7.6) overnight. To confirm the conformational states of the CCMV control samples, the putative contracted (pH 4.6) and expanded (pH 7.6) forms were prepared separately by manually applying each sample to the grid and blotting for five seconds, followed by retraction of the blotting assembly and plunging of the grid. Our ~3.2 Å reconstruction of the low pH CCMV (**Fig S5, Fig S6, Table S1**) and our ~6.9 Å reconstruction of the high pH CCMV (**Fig S7, Table S1**) are consistent with the contracted and expanded states previously described^17,29^, exhibiting diameters of 29 nm and 32 nm, respectively (**Fig 3C**).

**Figure 3.**
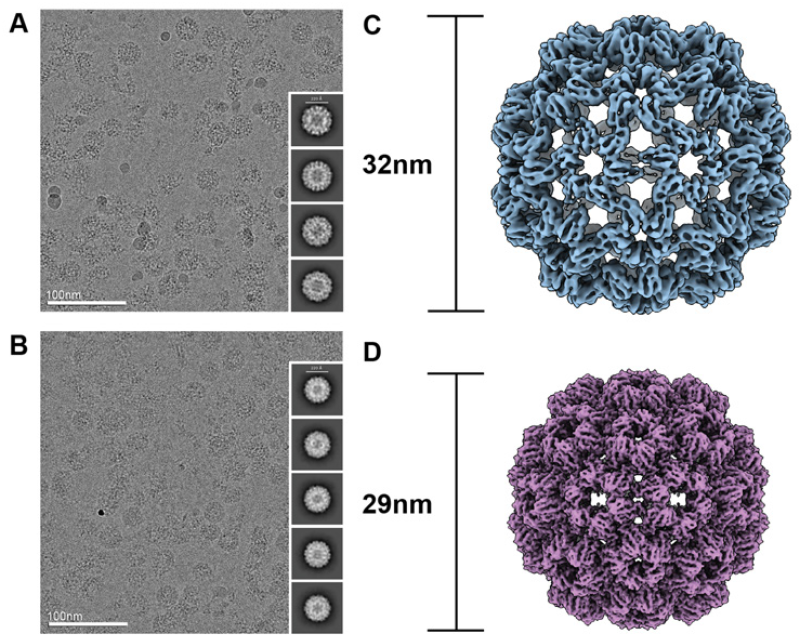
pH-induced contraction of CCMV using MIU. **A, B**) Representative micrographs and 2D class averages of the expanded state control (A) used as a starting point for the pH-shift experiment and the state resulting from contraction induced by application of low pH buffer with MIU (B). **C, D)** Representative maps of the expanded state control with a diameter of 32 nm (C) and the pH-shift-induced contracted state at 29 nm (D).

To determine if capsid contraction could be triggered through sample preparation with MIU, we first applied expanded CCMV sample at pH 7.6 to the grid and blotted for five seconds. The filter paper was held in the blotting position as low pH buffer (pH 3.6) mixed with gold fiducials was sprayed for 300 or 700 ms onto the opposite side of the grid to promote on-grid mixing of the capsids and buffer. Immediately after spraying, the filter paper was retracted, and the grid was plunged into the cryogen CCMV capsids were observed in the presence of gold fiducials throughout the acquired micrographs, confirming mixing of the capsids and the low pH buffer. Most micrographs containing capsids also contained 1-3 nanogold fiducials, though the local concentration varied, with clusters of more than 15 beads sometimes present in the micrographs. Capsids were also observed infrequently in micrographs without fiducials, though the neighboring micrographs contained gold, indicating some variability in the local concentration of fiducials. We observed a similar level of variation in the local distribution of fiducials during initial testing of a grid prepared with buffer as the primary sample and sprayed with a mixture of apoferritin and nanogold fiducials (data not shown). In this prior dataset, nanogold was observed in the majority of the images containing containing apoferritin, with only a few instances of apoferritin appearing in the absence of nanogold; thus, we concluded that the fiducials serve as a good approximation of the location of sprayed sample on the grid.

In the CCMV dataset, many regions of denatured capsids were observed, which was consistent with the data presented in the melting/re-vitrification CCMV pH shift study^17^, possibly indicating the structural instability of the capsid under such low pH conditions. Our ~3.9 Å reconstruction corresponded to a fully contracted state, with a diameter of 29 nm (**Fig 3D, Fig S8, Table S1**), consistent with the expected conformational state of CCMV capsids under low pH conditions^17,29^.

### MIU captures ATP-dependent GroEL/ES complex formation

Ligand-mediated protein conformational changes are attractive targets for time-resolved studies of biological systems due to rapid diffusion of small molecules. However, capturing these events remains technically challenging, with only a few successful examples demonstrated using highly specialized crystallographic techniques, such as serial femtosecond crystallography^30–33^. These methods often require extensive optimization and incur high costs for facility usage. Time-resolved cryo-EM offers a more versatile approach, enabling structural studies of larger and more complex proteins in a near-native state while accommodating a wider range of conformational dynamics. Further, the development of custom-built, cost-effective instrumentation for rapid sample preparation substantially reduces experiment-associated costs.

Several time-resolved cryo-EM studies have used ligands or small substrates to probe rapid conformational changes such as ATP-dependent dissociation of actomyosin^28^, ss-DNA-RecA filament growth^19^, and GTP-dependent dynamin conformational rearrangements^16^. We therefore investigated the capacity of rapid on-grid mixing using MIU to capture the assembly of the bacterial chaperonin GroEL/ES complex. The GroEL/ES machinery has served as a model system for understanding ATP-dependent protein folding and as a benchmark for studying protein complex assembly by cryo-EM^34–36^. GroEL forms a homo-tetradecameric assembly in the absence of GroES and ATP. Upon ATP binding, the apical heptamer undergoes a conformational change that facilitates GroES binding, thereby capping the complex and creating an enclosed cavity where unfolded or misfolded substrates can refold^37,38^. GroEL/ES capping has previously been observed to occur on the millisecond to second timescale^36^.

To establish the baseline organizational states of GroEL/ES in the absence of ATP, a mixture of GroEL/ES was prepared and incubated at room temperature for ten minutes prior to vitrification. The sample was manually applied to the grid, blotted, and vitrified, without addition of a sprayed sample. Uncapped GroEL, corresponding to a “barrel” conformation, was present uniformly in the data. Our ~3.3 Å map shows a lack of nucleotide density in the nucleotide binding pocket (**Fig 4A, Fig S9, Table S2**). GroES was also present within the sample and observed in 2D class averages, often forming a hexagonally-packed lattice with strong preferred orientation (**Fig S9**). Extensive rounds of 2D classification were performed to confirm the absence of capped GroEL particles.

**Figure 4.**
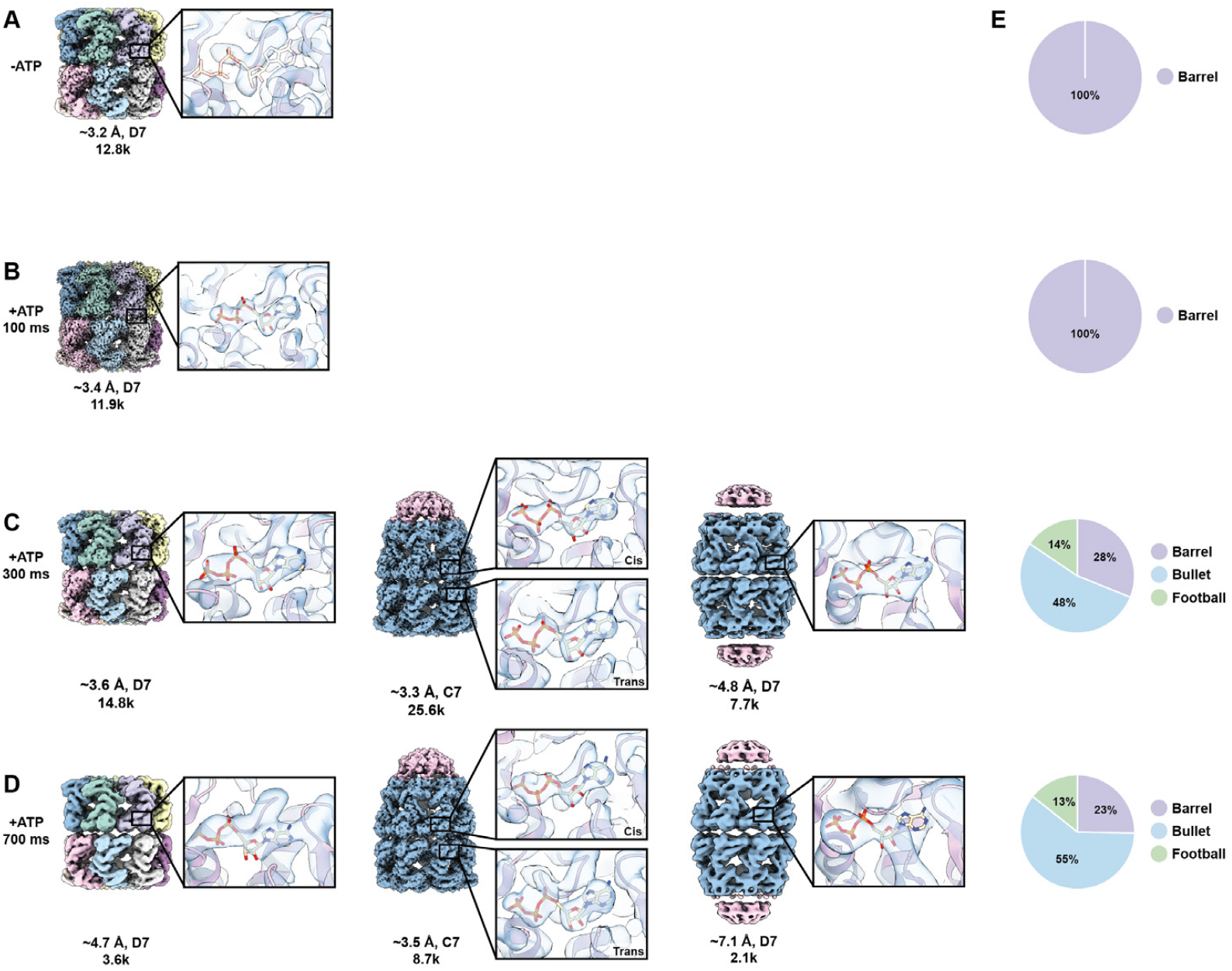
ATP-dependent complex formation of GroEL/ES observed following sample preparation with MIU. Density maps obtained from preparation of: **A)** the control GroEL in the presence of GroES without ATP; **B)** GroEL in the presence of GroES and sprayed ATP after 100 ms; **C)** GroEL + GroES with sprayed ATP after 300 ms; **D)** GroEL + GroES with sprayed ATP after 700 ms. The insets display representative subunit nucleotide density. **E)** Particle distributions for the control, 100 ms, 300 ms, and 700 ms from top to bottom, respectively.

We next assessed whether the blot-then-spray approach could drive GroEL/ES complex formation upon on-grid mixing. GroEL/ES samples were prepared as described for the ATP-free condition, with the addition of sprayed ATP supplemented with nanogold fiducials to identify regions of sprayed ATP localized on the grid. At the 100 ms timepoint, we were unable to detect GroEL/ES assembly, as all the GroEL particles were observed in the “barrel” conformation, despite performing several rounds of 2D classification in an attempt to identify a subpopulation of GroEL/ES particles (**Fig 4B, Fig S10**). Despite remaining in an uncomplexed state, nucleotide density was observed in the nucleotide binding pocket corresponding to an ATP molecule, indicating that 100ms was sufficient for nucleotide-binding (**Fig 4B**). This data is consistent with FRET data demonstrating ATP-binding to GroEL within 10ms^39^, as well as with a previous time-resolved cryo-EM study of GroEL/ES complex formation^36^. Although GroEL was observed complexed with GroES in the previous time-resolved study after 50ms, the proportion of these complexes was low (~4.2%). This discrepancy may be due to the difference in on-grid mixing behavior of particles in our study relative to the microfluidics system used in the former study.

At both the 300 ms and 700 ms timepoints, the majority of the GroEL particles were observed complexed with GroES, predominantly in the single-capped “bullet” conformation, with one GroES heptamer bound to one GroEL ring. Multiple rounds of 2D classification further revealed the presence of a double-capped GroEL/ES “football” complex in which GroES caps both ends of the GroEL barrel. 2D classification also revealed a small population of uncapped GroEL barrels. At the 300 ms timepoint, 48% of the extracted particles adopted the bullet conformation, 28% exhibited the barrel conformation, and 14% were in the football conformation (**Fig 4C, Fig S11**). These proportions were similar in the 700 ms dataset, with 55%, 23%, and 13%, corresponding to the bullet, barrel, and football assemblies, respectively (**Fig 4D, Fig S12**), suggesting that the GroEL/ES assembly reaction had largely reached equilibrium within the first 300 ms following ATP addition.

All three conformations exhibited nucleotide density in the nucleotide-binding pockets, including the uncapped GroEL barrels in both datasets (**Fig 4C,D**). The density was unambiguously resolved as ATP, except in the cases of the 700 ms barrel conformation and both the 300 ms and 700 ms football conformations, where low particle counts limited our attainable resolution, which prevented confident assignment of nucleotide state. These findings are consistent with presteady-state kinetic data showing that ATP hydrolysis occurs on the second timescale^40^. Interestingly, the barrel conformation of the 300 ms and 700 ms datasets closely resemble the GroEL conformation observed in the previous time-resolved study (PDB:8BL7), in which the apical domains of both heptameric rings are angled downward, reflecting an ADP-like state, despite exhibiting density for ATP in the nucleotide pockets. This may reflect a transient state between nucleotide binding and the rearrangement of the apical domains to accommodate GroES binding. The bullet conformation of these datasets exhibits a similar ADP-like conformation in the trans ring of GroEL and ATP-like conformation of the cis ring, despite density for ATP observed in both rings, reflecting a state that closely resembles another conformation observed in the previous time-resolved study (PDB:8BMO).

Taken together, these data confirm the ability of MIU to capture ligand-induced conformational states in protein complexes through rapid on-grid mixing.

## Discussion

In this study, we present Mix-it-up (MIU) - an updated and low-cost methodology for on-grid sample mixing, enabling the study of conformational changes in biological systems on the millisecond timescale. We first demonstrate that MIU improves the overall quality and consistency of ice on the grid suitable for imaging compared to previous iterations of the device (Shake-it-off, Back-it-up)^22,23^. To illustrate the feasibility of on-grid mixing, we used MIU to combine distinct protein samples - manually applied aldolase and sprayed apoferritin – serving as a proof of principle. We further show that this system supports the study of environmentally or ligand-induced conformational changes in protein samples with CCMV and the GroEL/ES chaperonin complex.

We also demonstrate the use of nanogold fiducials to localize spray distribution and localize regions of sample mixing on the grid. While we used fiducials in buffer or mixed with ligand solutions, this strategy could be extended to protein samples to track the spatial distribution of sprayed proteins that are too small or disordered to resolve by cryo-EM in an un-complexed form.

Despite the accessibility and cost-effectiveness of this MIU setup, we acknowledge that achieving full automation and greater reproducibility will require further engineering advances to improve precision and timing control. Further, we acknowledge that MIU does not currently achieve the temporal resolution required to capture conformational changes occurring on the microsecond or single-millisecond timescale. However, it represents a substantial improvement over traditional blotting-based methods, which typically require several seconds for sample preparation. By enabling sample mixing and vitrification within a few hundred milliseconds, MIU opens new possibilities for examining dynamic biological systems that evolve on intermediate timescales. As such, MIU offers a robust and adaptable platform for investigating transient states in macromolecular assemblies by single-particle cryo-EM, and lays the groundwork for future time-resolved cryo-EM studies of rapid biological processes, such as catalysis, signaling, and transport.

## Materials and Methods

### MIU build

MIU was initially assembled following instructions for the Shake-it-up and BIU devices designed by the Rubinstein group, as previously described^22,23^. Several modifications were made to the device, which included a redesign of the 3D-printed tweezer-to-solenoid connection, the blotting clip, and the blotting mount, as shown in (**Fig S1**). The STL files of these redesigned components are available at https://github.com/laurenalex77/Mixit-up. The software repository was downloaded from https://github.com/johnrubinstein/BIUcontrol.git and modified as follows: A method to enable the blot-spray-and-plunge sequence was added to the BIUapplyandplunge.py script and BIUgui.py, along with a parameter to specify the blot time within the GUI. A 60 second delay was also added following the plunge instruction in BIUapplyandplunge.py to enable adequate time for disconnection of the tweezers prior to transferring the grid to liquid nitrogen from the ethane bath.

### Sample preparation

#### Apoferritin

Heavy-chain mouse apoferritin was purified from frozen cell pellets prepared as previously described^41^. A pET24a vector was gifted from M. Kikkawa (University of Tokyo) and was transformed into Rosetta *E. coli* cells. Cells were grown at 37 °C in LB media and protein expression was induced with 1mM IPTG at OD_600nm_ = 0.5. Protein was expressed for 3 hours at 37 °C, after which cells were harvested via centrifugation and stored at −80 °C. Cell pellets were thawed and resuspended in lysis buffer containing 30 mM HEPES pH 7.5, 300 mM NaCl, 1 mM MgSO4, 1x Roche inhibitor, 30 mg lysozyme and lysed by sonication for a total of 10 min at 30% amplification with a pulse of 3s on / 8s off. The lysate was clarified via centrifugation at 20,000 RCF, 4 °C for 20 min. Heat precipitation was performed to remove host cell proteins by incubating the clarified lysate in a 70 °C water bath for 10 min and performing centrifugation at 20,000 RCF, 4 °C for 15 min. Ammonium sulfate precipitation was performed by adding 60% (w/v) ammonium sulfate to the supernatant, stirring on ice for 10 min, and centrifuging at 14,000 RCF, 4 °C for 20 min. The pellet containing apoferritin was resuspended in PBS and dialyzed in a 10 kDa MW cut-off dialysis cassette overnight in dialysis buffer (30 mM HEPES pH 7.5, 20 mM NaCl, 1 mM DTT) to remove ammonium sulfate. Ion exchange was performed the next day using two connected 5 mL Hi Trap Q HP columns. The column was washed with low salt buffer (30 mM HEPES pH 7.5, 20 mM NaCl, 1 mM DTT) and eluted from a gradient of 0-100% high salt buffer (30 mM HEPES pH 7.5, 500 mM NaCl, 1 mM DTT). Fractions containing apoferritin were pooled and concentrated using a 100 kDa MW concentrator. Apoferritin was further purified by size exclusion chromatography (SEC) using a Superdex 200 10/300 GL (Cytiva) column and SEC buffer (30mM HEPES pH 7.5, 150 mM NaCl, 1 mM DTT). Fractions containing apoferritin were pooled, concentrated to 14 mg/mL, and flash-frozen in 30 mM HEPES pH 7.5 + 150 mM NaCl.

#### Aldolase

Lyophilized rabbit muscle aldolase (Sigma-Aldrich) was solubilized in 20 mM HEPES pH 7.5, 50 mM NaCl to a final concentration of 1.4 mg/mL.

#### Cowpea Chlorotic Mottle Virus

CCMV was obtained as a gift from the Nicole Steinmetz lab at the University of California, San Diego. Preparation of CCMV was performed as previously described^42^. CCMV was harvested 14 days after inoculation of *Vigna unguiculata*, California black-eyed pea No. 5 plants. CCMV was purified following homogenization of leaves in buffer consisting of 0.2 M NaOAc and 1 mM EDTA at pH 4.8. The homogenate was filtered through cheesecloth, centrifuged, and the supernatant was precipitated overnight with 0.02 M NaCl and 8% (w/v) PEG. Following centrifugation, the pellet was resuspended in the contracted state buffer consisting of 0.1 M NaOAc and 1 mM EDTA at pH 4.8. The sample was subjected to further centrifugation and the supernatant was applied to a 20% (w/v) sucrose cushion and centrifuged for an additional 2 hr at 4 °C, 148,000 xg. The pellet was resuspended in the contracted state buffer and stored as an intact virion at −20°C until further use.

#### GroEL and GroES

*E. coli* GroEL and Gro ES were received as a gift from the Steven Johnson lab at the University of Indiana. Protein expression and purification of GroEL and GroES were performed as previously described^43^. T7 Express *E. coli* (cam^R^) containing a *Trc*-promoted GroES/GroEL plasmid (amp^R^) were cultured in LB media and induced with 1mM IPTG at OD_600_ ^≈^ 0.8. Cells were lysed by microfluidization after resuspension of cell pellets in 50 mM Tris pH 7.4, 50 mM KCl, 1 mM DTT, and 1 mM PMSF. The lysed cells were clarified by centrifugation (22k xg for 45min), and the lysate was purified via cation exchange using FFQ (Cytiva) resin in buffer containing 50 mM Tris pH 7.4, 1 mM DTT, with a 0 to 1 M NaCl gradient. GroEL-containing fractions were precipitated with 1.2 M ammonium sulfate at 4°C and further purified via hydrophobic interaction chromatography using source 15ISO resin (Cytiva) in buffer containing 50 mM Tris pH 7.4, 50 mM KCl, 150 mM NaCl, and 1 mM DTT. GroEL fractions were then dialyzed into storage buffer containing 50 mM Tris pH 7.4, 150 mM NaCl, and 1 mM DTT.

*E. coli G*roES was similarly purified with the following exceptions: after cation exchange, Gro-ES containing fractions were adjusted to pH 4.6 with 50mM sodium acetate buffer. The sample then further purified by additional cation exchange using a FFSP (Cytiva) column in buffer containing 50 mM Tris pH 4.6, 1 mM DTT, with a 0 to 1 M NaCl gradient. GroES-containing fractions were size-exchanged on a Hiload 26/600 Superdex 75 (GE) column and eluted in 50 mM Tris pH 7.4, 300 mM NaCl, and 1 mM DTT. Fractions were concentrated GroES were concentrated and stored.

GroEL and GroES stocks used for cryo-EM sample preparation were diluted as needed in buffer containing 50 mM Tris, pH 7.4, 50 mM KCl, 1 mM MgCl_2_.

#### Gold fiducials

Ted Pella 5nm gold colloid was prepared according to Iancu et al., 2006^44^ (section “First Phase”, Tube B). 5x the desired sample volume of fiducials was centrifuged at 18,000 xg for 25 min at 4°C. The pellet containing the fiducials was resuspended in sample, resulting in a 5x final concentration of fiducials.

### Cryo-EM grid preparation

The general procedure for grid preparation is as follows: A homemade, holey gold 2.2/2.2, 400 EM grid prepared as previously described^45^ or an UltrAuFoil R2/2, 200 was plasma cleaned for seven seconds at 25 Watts (75% argon, 25% oxygen) using a Solarus plasma cleaner (Gatan, Inc.).

For single-sample experiments, only the sample side of the grid was plasma cleaned. Both sides were cleaned for spray experiments. The grid was then mounted to the tweezers above the cryogen bath with the smooth side facing towards the blotting paper and the back side (copper) positioned in front of the piezo. A 0.5 × 2 cm rectangle of Whatman No.1 filter paper was positioned in the blotting clip in front of a similarly sized strip of cellulose acetate. 3 µL of sample A was manually applied to the smooth side of the grid. 2 µL of sample B was applied to the back side of the piezo (opposite the side facing the grid). The spray time, retraction delay, and plunge delay were set to 100, 300, or 700 ms, and the blot time was set to 5 seconds on the MIU GUI. The “Blot, Spray, and Plunge” button was pressed to trigger 5 seconds of blotting, followed by the specified spray time, filter paper retraction, and plunging into liquid ethane.

#### Aldolase + apoferritin

Sample A, 7 mg/mL of apoferritin in 30 mM HEPES pH 7.5 + 150 mM NaCl, was manually applied to a homemade grid. Sample B, 1.4 mg/mL of aldolase in 20 mM HEPES pH 7.5, + 50 mM NaCl, was applied to the piezo.

#### Contracted CCMV control

Sample A, 10 mg/mL of contracted CCMV in 0.1 M NaOAc, 1 mM EDTA, pH 4.8, was manually applied to a homemade grid.

#### Expanded CCMV control

Sample A, 20 mg/mL of expanded CCMV in 10mM Tris (base), 50mM NaCl, 1mM EDTA, pH 7.6, was manually applied to a homemade grid.

#### Expanded CCMV + low pH buffer / gold (700 ms)

Sample A, 20 mg/mL expanded CCMV in 10mM Tris (base), 50mM NaCl, 1mM EDTA, pH 7.6, was manually applied to an UltrAuFoil 2/2, 200 grid. Sample B, low pH buffer consisting of 0.1M NaOAc, 1mM EDTA at pH 3.6, was applied to the piezo.

#### GroEL + GroES, no ATP

Sample A, 8.6 µM of GroEL + 17.1 of GroES, was incubated at room temperature for 10-15min and applied manually to the grid. Spray time, plunge delay, and retraction delay were all set to 700 ms.

#### GroEL + GroES + ATP/gold (100 ms)

Sample A, 8.6 µM GroEL + 17.1 µM GroES, was incubated at room temperature for 10-15 min prior to freezing and applied manually to a homemade grid. Sample B, 83.3 mM ATP, was applied to the piezo. Spray time, plunge delay, and retraction delay were all set to 100 ms.

#### GroEL + GroES + ATP/gold (300 ms)

Sample A, 8.6 µM of GroEL + 17.1 µM of GroES, was the same mixture used for the no-ATP-control and was applied manually to an UltrAuFoil 2/2 grid following incubation at RT. Sample B, 83.3 mM ATP, was applied to the piezo. Spray time, plunge delay, and retraction delay were all set to 300 ms.

#### GroEL + GroES + ATP/gold (700 ms)

Sample A, 8.6 µM of GroEL + 17.1 µM of GroES, was the same mixture used for the no-ATP-control and was applied manually to an UltrAuFoil 2/2 grid following incubation at RT. Sample B, 83.3 mM ATP, was applied to the piezo. Spray time, plunge delay, and retraction delay were all set to 700 ms.

The plunge delay for MIU experiments is approximately 23.1 ms. This average was calculated based on three separate videos from experiments where the spray time was set to 100 ms, 300 ms, and 700 ms. The number of frames between the initiation of spray and the submergence of the grid in liquid ethane were counted. There were 24 frames in the 100 ms experiment video (200.58 fps), 64 frames in the 300 ms experiment video (200.43 fps), and 147 frames in the 700 ms experiment video (201.21 fps). Subtracting the time of spray, the resulting plunge delays were calculated to be 19.6 ms, 19.3 ms, and 30.4 ms, yielding an average of 23.1 ms.

### Data collection

Cryo-EM data was collected on a Talos Arctica equipped with a 200 kV field emission gun and a Falcon 4i direct electron detector. Micrographs were collected at 150,000 x nominal magnification (yielding a pixel size of 0.94 Å /pixel at the detector) using the EPU data collection software. The specified defocus range was −1.4, −1.2, −1 µm. The mixed apoferritin/aldolase dataset was collected at an exposure rate of 10.53 e^−^/pixel/s with an exposure time of 4.22 s.

The contracted CCMV control dataset was collected at an exposure rate of 10.75 e^−^/pixel/s with an exposure time of 4.13 s. The expanded CCMV control dataset was collected at an exposure rate of 9.93 e^−^/pixel/s with an exposure time of 4.48 s. The CCMV pH shift dataset was collected at an exposure rate of 10.9 e^−^/pixel/s with an exposure time of 4.08 s. The GroEL/ES, no ATP and +ATP 700 ms datasets were collected at an exposure rate of 10.9 e^−^/pixel/s with an exposure time of 4.08 s. The GroEL/ES + ATP 300 ms dataset was collected at an exposure rate of 9.66 e^−^/pixel/s with an exposure time of 4.61 s. The GroEL/ES + ATP 100 ms dataset was collected at an exposure rate of 10.94 e^−^/pixel/s with an exposure time of 4.08 s.

#### Aldolase + apoferritin

The image processing pipeline is shown in (**Fig S4**). 2,476 movies were collected. Motion correction and CTF correction were performed in cryoSPARC Live v4.6.2. After applying a resolution cutoff of 6 Å based on CTF estimation, 1,673 micrographs remained for further processing. Apoferritin particles were picked with cryoSPARC live Blob Picker with a particle diameter of 100-110 Å, a NC score *>*0.3 and a power score range of −17 to 1,329. 1,071,352 particles were extracted with a box size of 200 px, binned to 100 px. 2D classification was performed with 50 classes. Subset selection was performed with stringent cleanup (the single class having the most well-defined secondary structure was selected) and minimal cleanup (only one class that did not have features consistent with apoferritin was excluded) in parallel, with 81,562 particles selected in the stringent instance and 1,049,548 selected in the minimal instance. *Ab initio* reconstruction with one class, octahedral symmetry, and the initial resolution set to 6 Å was performed on the stringent cleanup particles, followed by non-uniform refinement with octahedral symmetry, the initial lowpass resolution set to 8 Å, and per-particle defocus and per-group CTF parameter optimization turned on. *Ab initio* reconstruction with three classes and octahedral symmetry, and the initial resolution set to 8 Å was performed on the minimal cleanup particles. The reconstruction from non-uniform refinement of the stringent clean up particles was input into a heterogenous refinement job, along with two classes from the minimal clean up ab-initio to serve as junk classes for sorting. All extracted particles were input, and the refinement was run with octahedral symmetry and hard classification enforced. 465k particles were sorted into the “good” class as the output of the heterogenous refinement job. These particles were re-extracted without binning and further refined using non-uniform refinement with octahedral symmetry, the initial lowpass resolution set to 8 Å, and per-particle defocus and per-group CTF parameter optimization turned on, yielding a ~2.9 Å map, based on a Fourier Shell Correlation (FSC) cutoff of 0.143 (**Table S1**).

Aldolase particles were picked with cryoSPARC Blob Picker with a particle diameter of 100-110 Å, an NC score *>*0.12 and a power score range of 409 to 658. 820,058 particles were extracted with a box size of 200 px, binned to 100 px, and 2D classification was performed with 50 classes. Subset selection was preformed to yield 389,144 particles. *Ab initio* reconstruction with 2 classes, C1 symmetry, and the initial resolution set to 8 Å was performed. The “good” reconstruction was then further refined with non-uniform refinement using D2 symmetry, the initial lowpass resolution set to 8 Å, and per-particle defocus and per-group CTF parameter optimization turned on. This reconstruction was input into a heterogenous refinement job, along with duplicates of the “junk” class yielded during *Ab initio* reconstruction. All extracted particles were input and the refinement was run with D2 symmetry. 428k particles were sorted into the “good” class as the output of the heterogenous refinement job. These particles were re-extracted without binning and further refined using non-uniform refinement with D2 symmetry, the initial lowpass resolution set to 8 Å, and hard classification enforced. Further exposure curation was performed to apply a defocus cutoff of 1.3µm, a CTF fit resolution cutoff of 3.5 Å, a relative ice thickness cutoff of 1.1, and a defocus tilt angle cutoff of 10 degrees. 622 exposures and 185,420 particles remained after exposure curation. A reconstruction with the resulting particles was further refined with non-uniform refinement with D2 symmetry, the initial lowpass resolution set to 6 Å, and per-particle defocus and per-group CTF parameter optimization turned on, yielding a ~2.9 Å map, based on a Fourier Shell Correlation (FSC) cutoff of 0.143 (**Table S1**).

#### Contracted CCMV

The image processing pipeline is shown in (**Fig S6**). 292 micrographs were collected. Motion correction and CTF correction were performed in cryoSPARC live v4.6.2. CCMV particles were initially picked with cryoSPARC live blob picker using a diameter of 100-500 Å. After initial 2D classification in cryoSPARC live, the best two 2Ds were used as templates for live template picker with the particle diameter set to 280 Å, the minimum separation distance set to 0.2 diameters, the NC score set to *>* 0.3, and the power score range of −86 to 2,775. 7,994 particles were extracted with a box size of 500 px, binned to 100 px, and 2D classification was performed with 50 classes and a circular mask of 280 Å. *Ab initio* reconstruction with 2 classes, C1 symmetry, and the initial resolution set to 20 Å was performed. The “good” class containing 5,397 particles was further refined using non-uniform refinement with icosahedral symmetry, the initial lowpass resolution set to 20 Å, and per-particle defocus and per-group CTF parameter optimization turned on. These particles were re-extracted without binning and further refined using non-uniform refinement with icosahedral symmetry, the initial lowpass resolution set to 20 Å, and per-particle defocus and per-group CTF parameter optimization turned on, yielding a ~3.2 Å map, based on a Fourier Shell Correlation (FSC) cutoff of 0.143 (**Table S1**).

#### Expanded CCMV

The image processing pipeline is shown in (**Fig S7**). 595 micrographs were collected. A resolution CTF cutoff of 6 Å was applied. 342 micrographs remained after preprocessing. Motion correction and CTF correction were performed in cryoSPARC live v4.6.2. CCMV particles were initially picked with cryoSPARC live blob picker using a diameter of 200-500 Å. After initial 2D classification in cryoSPARC live, the best four 2Ds were used as templates for live template picker with the particle diameter set to 310 Å, the minimum separation distance set to 0.2 diameters, the NC score set to *>* 0.257, and the power score range of 1,030 to 1,839. 62,017 particles were extracted with a box size of 560 px, binned to 100 px, and 2D classification was performed with 50 classes and a circular mask of 370 Å. *Ab initio* reconstruction on a selected subset of 2,269 particles with 2 classes, C1 symmetry, and the initial resolution set to 20 Å was performed. The “good” class containing 1,885 particles was further refined using non-uniform refinement with icosahedral symmetry. These particles were re-extracted without binning and further refined using non-uniform refinement with icosahedral symmetry to yield a ~6.9 Å map, based on a Fourier Shell Correlation (FSC) cutoff of 0.143 (**Table S1**).

#### Expanded CCMV + low pH buffer / gold (700 ms)

The image processing pipeline is shown in (**Fig S8**). 6,007 micrographs were collected. A resolution CTF cutoff of 12 Å was applied. 3,050 micrographs remained after preprocessing. Motion correction and CTF correction were performed in cryoSPARC live v4.6.2. CCMV particles were picked with cryoSPARC live blob picker using a diameter of 275-325 Å, an NC score *>*0.156 and a power score range of −942 to 2,434. 68,810 particles were extracted with a box size of 560 px, binned to 100 px, and 2D classification was performed with 50 classes and a circular mask of 390 Å. *Ab initio* reconstruction on a selected subset of 6,952 particles with 1 class, C1 symmetry, and the initial resolution set to 20 Å was performed. The reconstruction was further refined using non-uniform refinement with icosahedral symmetry and the initial lowpass resolution set to 15 Å. These particles were re-extracted without binning and further refined using non-uniform refinement with icosahedral symmetry, the initial lowpass resolution set to 6 Å, and per-particle defocus and per-group CTF parameter optimization turned on, yielding a ~3.9 Å map, based on a Fourier Shell Correlation (FSC) cutoff of 0.143 (**Table S1**).

#### GroEL + GroES, no ATP

The image processing pipeline is shown in (**Fig S9**). 601 micrographs were collected. A resolution CTF cutoff of 7.5 Å was applied. 376 micrographs remained after preprocessing. Motion correction and CTF correction were performed in cryoSPARC live v4.6.2. Particles were picked in cryoSPARC with Topaz using the ResNet16 pretrained dataset, an estimated particle diameter of 240 Å, and a particle threshold of −5. The picks were selected based on an NC score *>* −15.56 and a power score range of −30 to 143. 60,116 particles were extracted with a box size of 420 px, binned to 100 px, and initial 2D classification was performed with 50 classes. Extensive 2D classification was performed iteratively in parallel to confirm the absence of capped GroEL. Classes resembling GroEL from the initial 2D classification were selected (10,969 particles) and an *ab initio* reconstruction with 2 classes and C1 symmetry, and the initial resolution set to 15 Å was performed. Both classes were used as inputs for a heterogenous refinement job with 17,477 particles from a less stringent 2D class subset selection, the initial lowpass resolution set to 12 Å, and hard classification enforced. The 11,859 particles sorted into the “good” class were then re-extracted without binning and the reconstruction was further refined using non-uniform refinement with D7 symmetry, the initial lowpass resolution set to 6 Å, and per-particle defocus and per-group CTF parameter optimization turned on, yielding a ~3.2 Å map, based on a Fourier Shell Correlation (FSC) cutoff of 0.143 (**Table S2**).

#### GroEL + GroES + ATP/gold (100 ms)

The image processing pipeline is shown in (**Fig S10**). 2,645 micrographs were collected. A CTF cutoff of 6 Å, relative ice thickness cutoff of 1.1, total motion cutoff of 50 pixels, and defocus range cutoff of 110 Å was applied. 1,434 micrographs remained after preprocessing. Motion correction and CTF correction were performed in cryoSPARC live v4.6.2. Particles were picked in cryoSPARC with Topaz using the ResNet16 pretrained dataset, an estimated particle diameter of 240 Å, and a particle threshold of −5. The picks were selected based on an NC score *>* 0 and a power score of 25 – 120. 119,235 particles were extracted with a box size of 420 px, binned to 100 px, and initial 2D classification was performed with 50 classes. Classes resembling GroEL were selected for a total of 11,602 particles. Extensive 2D classification was performed iteratively to confirm the absence of capped GroEL. *Ab initio* reconstruction with C1 symmetry was performed with 1 and 2 classes and the initial resolution set to 15 Å in parallel. The reconstruction from the 1-class job, which corresponds to the single-capped GroEL in the “bullet” conformation, was further refined using non-uniform refinement with C7 symmetry and the initial lowpass resolution set to 20 Å. The reconstructions were further refined in independent non-uniform refinement jobs with D7 symmetry and the initial lowpass resolution set to 15 Å. The refined “best” class from the 2-class a*b initio* and refined class from the 1-class *ab initio* were used as inputs for heterogenous refinement (C1 symmetry, enforced hard classification, initial resolution 12 Å) containing 17,663 particles resulting from a less stringent 2D classification subset selection. After sorting, the 11.9k particles from the best reconstruction were re-extracted without binning and further refined using non-uniform refinement with optimized per-particle defocus and per-group CTF parameters, D7 symmetry and an initial lowpass resolutions of 6 Å. The resulting reconstruction was resolved at ~3.4 Å, based on a Fourier Shell Correlation (FSC) cutoff of 0.143 (**Table S2**).

#### GroEL + GroES + ATP/gold (300 ms)

The image processing pipeline is shown in (**Fig S11**). 5,301 micrographs were collected. A resolution CTF cutoff of 7.5 Å was applied. 3,665 micrographs remained after preprocessing. Motion correction and CTF correction were performed in cryoSPARC live v4.6.2. Particles were picked in cryoSPARC with Topaz using the ResNet16 pretrained dataset, an estimated particle diameter of 240 Å, and a particle threshold of −5. The picks were selected based on an NC score *>* 27.16 and a power score range of 30 to 149. 255,398 particles were extracted with a box size of 420 px, binned to 100 px, and initial 2D classification was performed with 50 classes. Classes resembling GroEL were selected for a total of 53,790 particles. *Ab initio* reconstruction with C1 symmetry was performed with 1 and 2 classes and the initial resolution set to 15 Å in parallel. The reconstruction from the 1-class job, which corresponds to the single-capped GroEL in the “bullet” conformation, was further refined using non-uniform refinement with C7 symmetry and the initial lowpass resolution set to 15 Å. Further rounds of 2D classification revealed a class corresponding to uncapped GroEL in the “barrel” conformation containing 2,230 particles, as well as a class corresponding to a double-capped GroEL in the “football” conformation with 761 particles. Both particle subsets were reconstructed independently using *ab initio* reconstruction with 1 class, C1 symmetry, and the initial resolution set to 20 Å. The reconstructions were further refined in independent non-uniform refinement jobs with D7 symmetry and the initial lowpass resolution set to 15 Å. The refined reconstructions of the bullet, barrel, and football conformations, along with a “junk” class from the initial 2-class *ab initio* job were used as inputs for heterogenous refinement (C7 symmetry, enforced hard classification, initial resolution 12 Å) containing the 53,790 particles resulting from the initial 2D classification subset selection. After sorting, 14.8k particles were assigned to the barrel conformation, 7.7k to the football, and 25.6k to the bullet. Particles from each conformation were re-extracted without binning and the reconstructions were further refined using non-uniform refinement with optimized per-particle defocus and per-group CTF parameters, using D7, D7, and C7 symmetry and initial lowpass resolutions of 6 Å, 8 Å, and 6 Å for the barrel, football, and bullet conformations, respectively.

The resulting reconstructions were resolved at ~3.6 Å for the barrel conformation, ~4.8 Å for the football conformation, and ~3.3 Å for the bullet conformation, based on a Fourier Shell Correlation (FSC) cutoff of 0.143 (**Table S2**).

#### GroEL + GroES + ATP/gold (700 ms)

The image processing pipeline is shown in (**Fig 12**). 6,063 micrographs were collected. A resolution CTF cutoff of 7.5 Å was applied. 2,535 micrographs remained after preprocessing. Motion correction and CTF correction were performed in cryoSPARC live v4.6.2. Particles were picked in cryoSPARC with Topaz using the ResNet16 pretrained dataset, an estimated particle diameter of 240 Å, and a particle threshold of −5. The picks were selected based on an NC score *>* 25.96 and a power score range of 47 to 139. 118,543 particles were extracted with a box size of 420 px, binned to 100 px, and initial 2D classification was performed with 50 classes. Classes resembling GroEL were selected for a total of 15,977 particles. The data was processed as described in the 300 ms dataset above with the following exceptions: *ab initio* reconstructions were performed with the initial resolution set to 20 Å, except for the bullet *ab initio* (15 Å). After sorting with heterogenous refinement, 3.6k particles were assigned to the barrel conformation, 2.1k to the football, and 8.8k to the bullet. Particles from each conformation were re-extracted without binning and further refined as described in the 300 ms dataset. The resulting reconstructions were resolved at ~4.7 Å for the barrel conformation, ~7.1 Å for the football conformation, and ~3.5 Å for the bullet conformation, based on a Fourier Shell Correlation (FSC) cutoff of 0.143 (**Table S2**).

## Acknowledgements

We thank Hannah Turner for help in assembling the initial BIU device, John Rubinstein (University of Toronto) and Jianhua Zhao (Sanford-Burnham-Prebys) for helpful discussions and advice on troubleshooting the ultrasonic spray device and grid fabrication. We also thank Mason Klawitter for help with grid fabrication, Tumara Boyd for SEM training on the Aquilos, the Danielle Grotjahn lab for providing the nanogold fiducials, Anthony Omole from the Nicole Steinmetz lab (UCSD) for CCMV samples, and the Steve Johnson lab (Indiana University School of Medicine) for GroEL/ES samples. LA is a Skaggs-Oxford Fellow and supported by an F31 Ruth L. Kirschstein Predoctoral Individual National Research Service Award from the National Institutes of Health (NIH), and support for these studies and GCL came from NIH grants GM143805 and GM154216.

## Data availability

All cryo-EM maps have been deposited in the Electron Microscopy Databank (EMDB) under the following accession codes: Apoferritin, EMD-70747; Aldolase, EMD-70748; CCMV contacted control, EMD-70749; CCMV expanded control, EMD-70750; CCMV pH shift, EMD-70751; GroEL control (no ATP), EMD-70752; GroEL “barrel”, 100 ms, EMD-70759; GroEL “barrel”, 300 ms, EMD-70753; GroEL “bullet”, 300 ms, EMD-70754; GroEL “football”, 300 ms, EMD-70755; GroEL “barrel”, 700 ms, EMD-70756; GroEL “bullet”, 700 ms, EMD-70757; GroEL “football”, 700 ms, EMD-70758. Updated 3D-printable components used in this paper can be found at https://github.com/laurenalex77/Mix-it-up.

## Supplementary Figures

**Figure S1.**
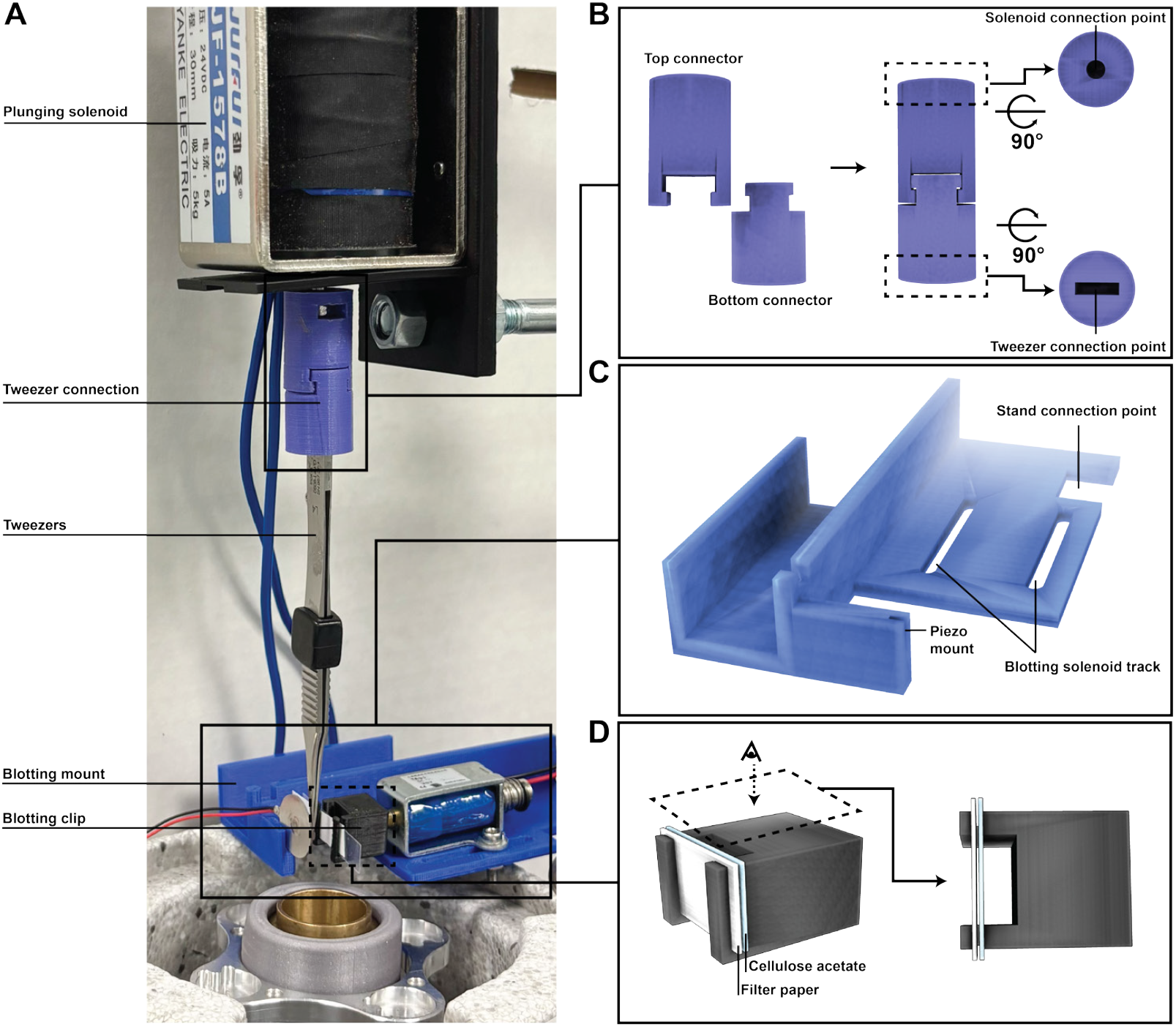
Modifications implemented in “Mix-it-up”. **A)** Photograph of the Mix-it-up device denoting modifications introduced. **B)** Schematic of the updated tweezer connection design, which consists of a top component that connects to the plunging solenoid and a bottom component that connects to the tweezers. The components are shown disjoined (left) and joined (right). The top and bottom views show the connection points to the solenoid and tweezers, respectively. **C)** Schematic of the blotting mount. **D)** Side (left) and top (right) views of the blotting clip with filter paper positioned between the two pillars and a strip of cellulose acetate used as backing support.

**Figure S2.**
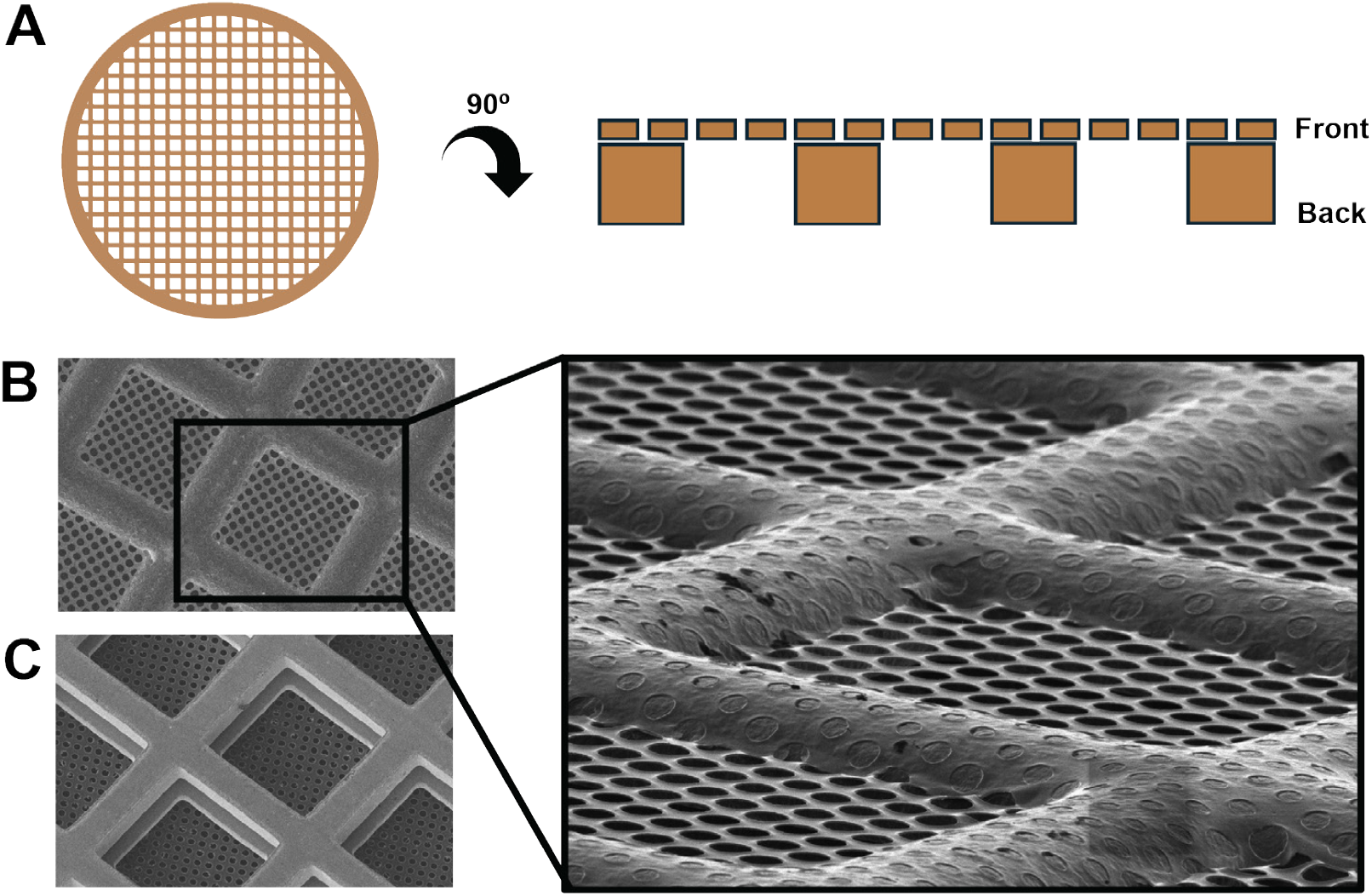
EM grid architecture. **A**) 2D schematic of an EM grid (left) and a representative schematic of the side view of several squares depicting the holey foil (front) and grid bars (back). **B)** Scanning electron microscopy (SEM) image of the front of a grid, showing a rounded surface of the grid bars overlaid with a holey foil (inset). **C)** SEM image of the back of a grid.

**Figure S3.**
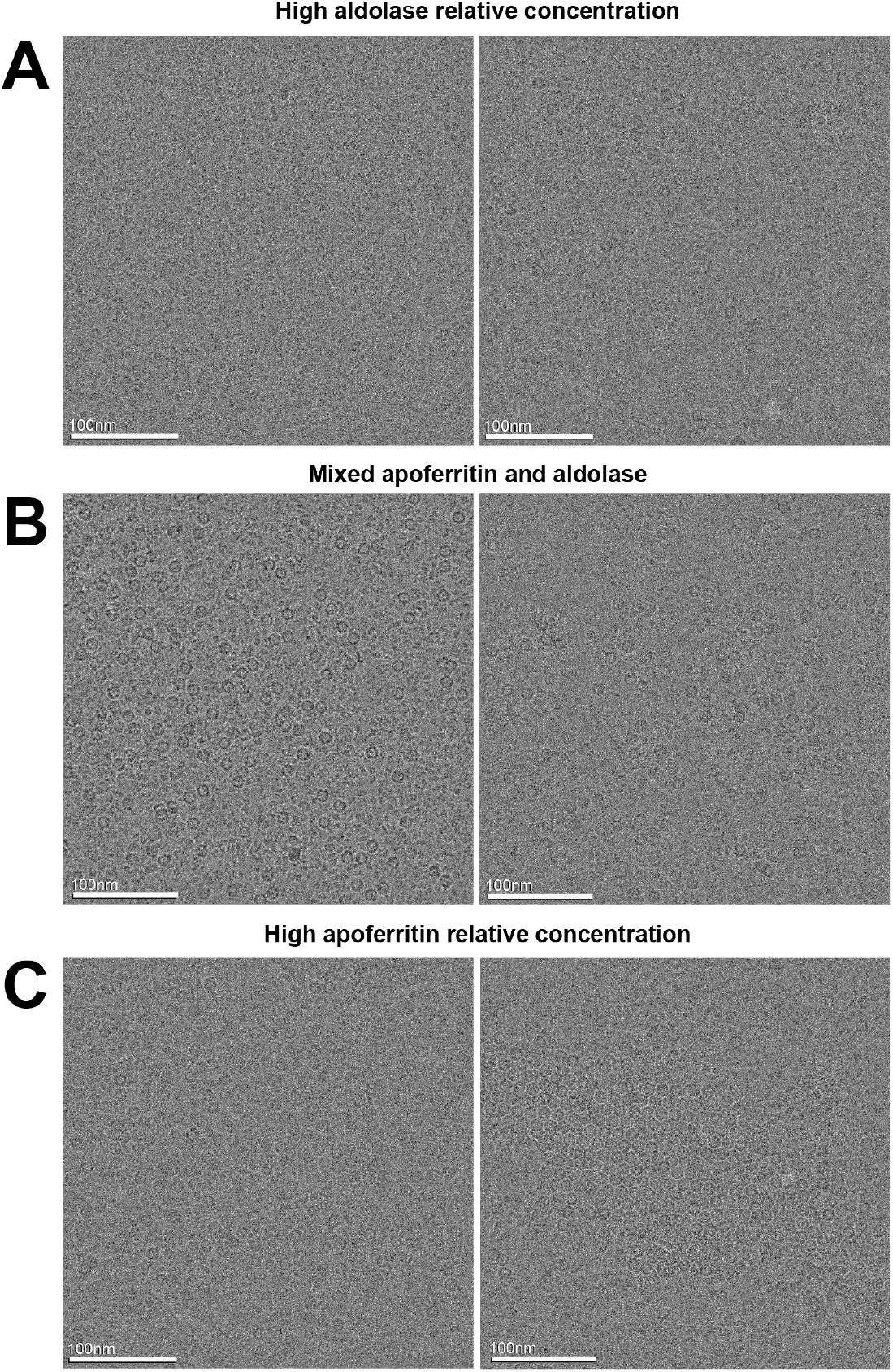
Aldolase and apoferritin particle distribution in mixed sample prepared with MIU. Representative micrographs from grids prepared with: **A)** high aldolase concentration, **B)** mixed apoferritin and aldolase concentrations, and **C)** high apoferritin concentrations.

**Figure S4.**
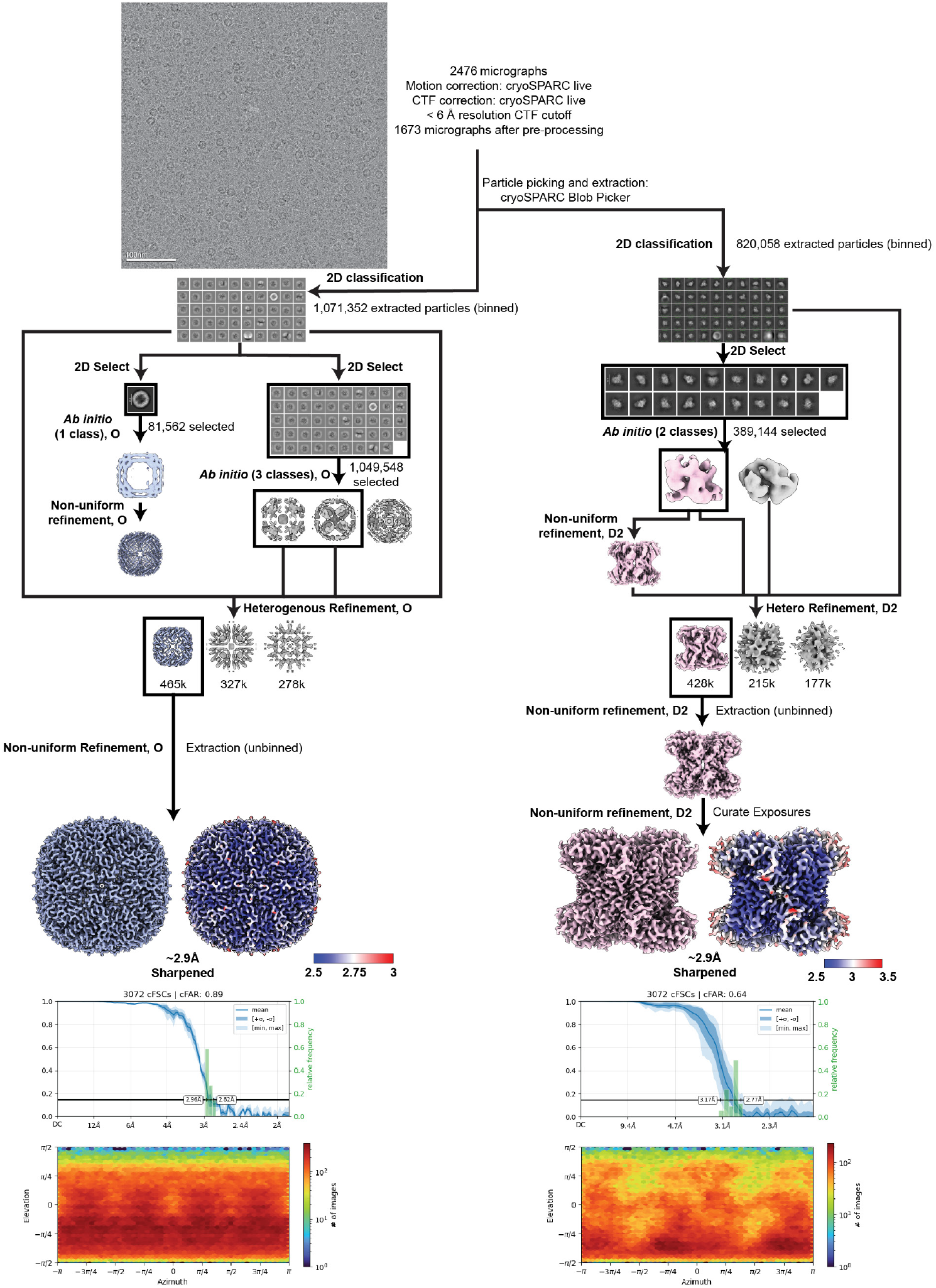
Processing pipeline of the mixed apoferritin and aldolase dataset prepared with MIU. Representative processing pipeline for mixed sample of apoferritin and aldolase. Motion and CTF correction were performed in cryoSPARC Live, as well as picking and extraction of apoferritin particles. Micrographs were exported to cryoSPARC for aldolase particle picking and extraction. Following 2D classification of apoferritin particles, two subsets were generated using stringent and minimal cleanup strategies and *ab initio* reconstructions were generated with octahedral symmetry imposed. The highest resolution *ab initio* class was used for initial 3D refinement with imposed symmetry. All extracted particles were then combined with several “good” and “junk” initial volumes in a heterogeneous refinement job with symmetry imposed to separate high-quality particles. Aldolase particles were processed in a similar manner with the following exceptions: a single subset of particles was generated of the best class averages following 2D classification. Following *ab initio* reconstruction, initial refinement, and heterogenous refinement, the particles were further refined, and exposures were curated to further filter out poor quality particles. For both apoferritin and aldolase, final refinement of the selected particles yielded a map at ~2.9 Å resolution. Final maps are shown next to a local resolution map. The conical FSC plot overlaid with a resolution histogram and corresponding angular distribution plot are shown below the final maps.

**Figure S5.**
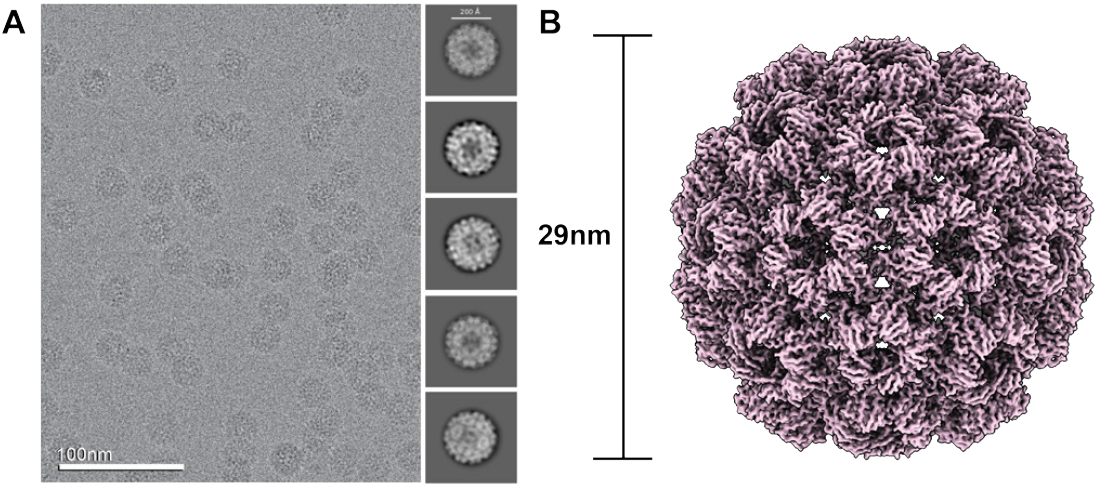
Control, contracted CCMV. **A)** Representative micrograph and 2D class averages of CCMV sample in pH 4.6 buffer. **B)** EM density map of the contracted state exhibiting a 29 nm diameter.

**Figure S6.**
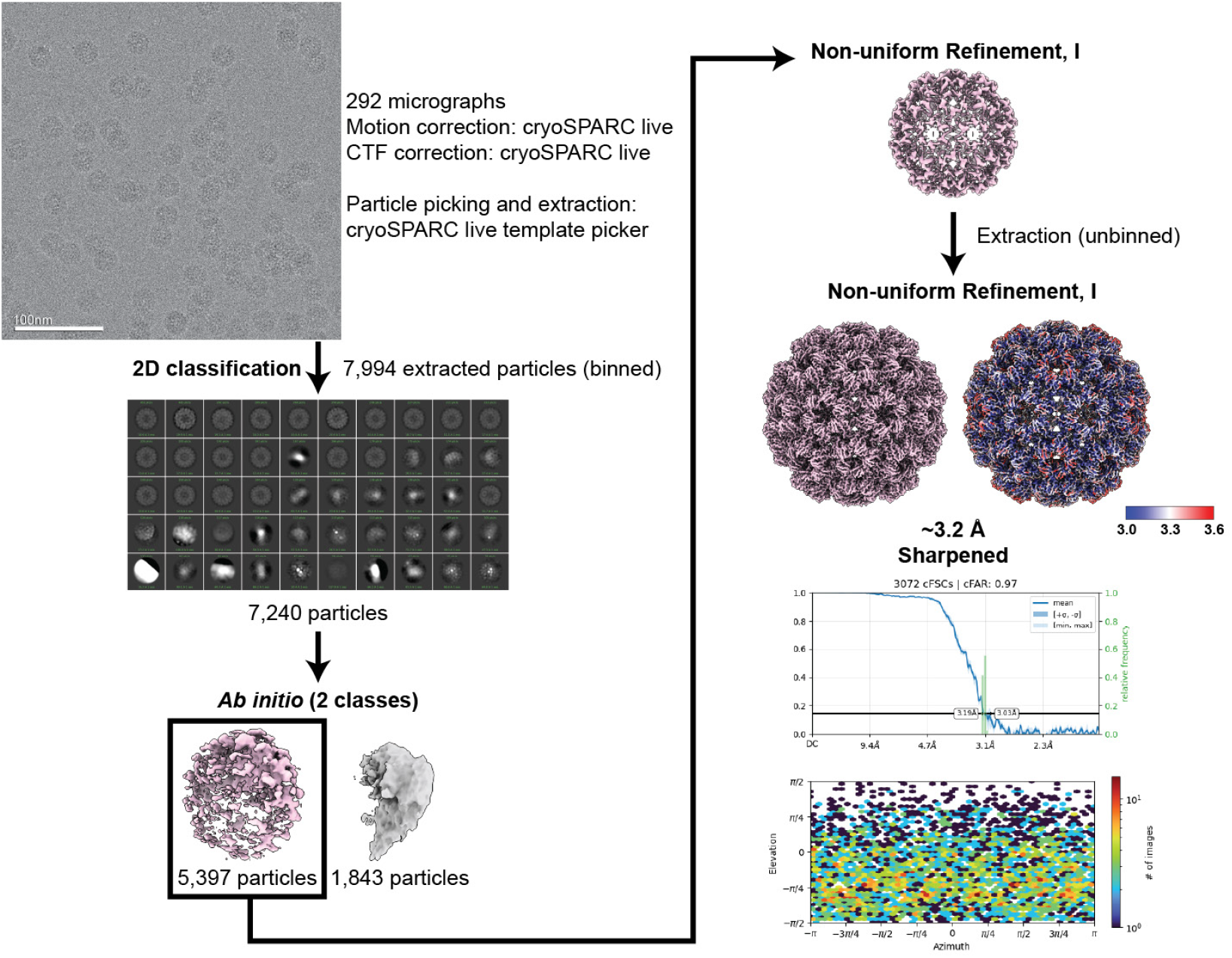
Processing pipeline of the control, contracted CCMV state dataset. Representative processing pipeline for CCMV dataset. Motion and CTF correction, initial particle picking, and initial 2D classification were performed in cryoSPARC Live. Template-based picking was used to refine particle selection, followed by additional 2D classification and *ab initio* reconstruction. Particles from the best class were used for 3D refinement with imposed icosahedral symmetry. After re-extraction, particles were further refined to yield a final map at ~3.2 Å resolution. The final map is shown next to a local resolution map. A conical FSC plot overlaid with a histogram of resolutions and an angular distribution plot are shown below the final maps.

**Figure S7.**
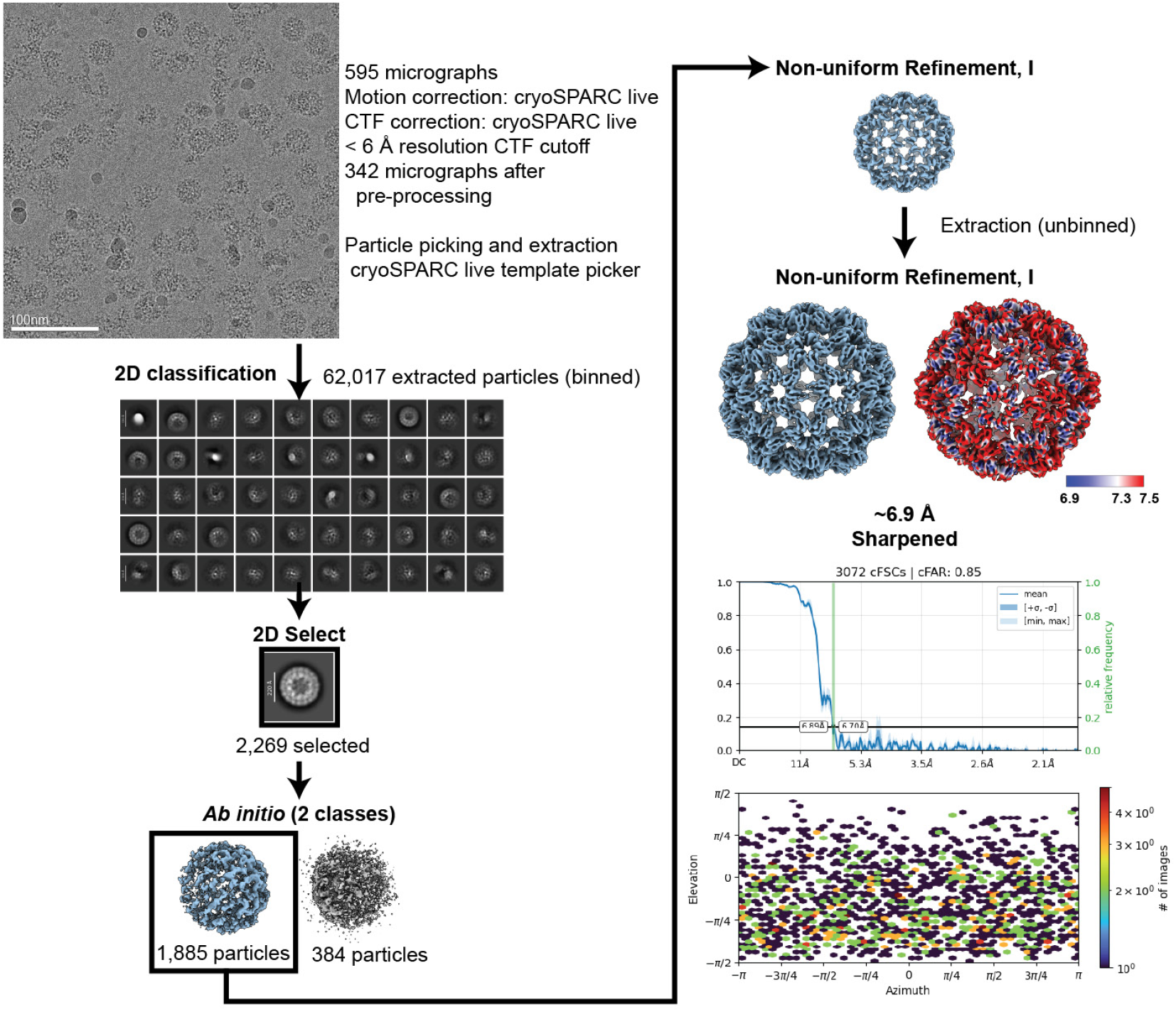
Processing pipeline of the control, expanded CCMV state dataset. Motion and CTF correction were performed in cryoSPARC Live, with a resolution cutoff applied during preprocessing. Initial blob-based particle picking was followed by template-based picking and 2D classification. A selected subset of particles was used for *ab initio* reconstruction and 3D refinement was performed with imposed icosahedral symmetry. After re-extraction, particles were further refined to yield a final map at ~6.9 Å resolution. The final map is shown next to a local resolution map. The conical FSC plot overlaid with a resolution histogram and corresponding angular distribution plot are shown below the final map.

**Figure S8.**
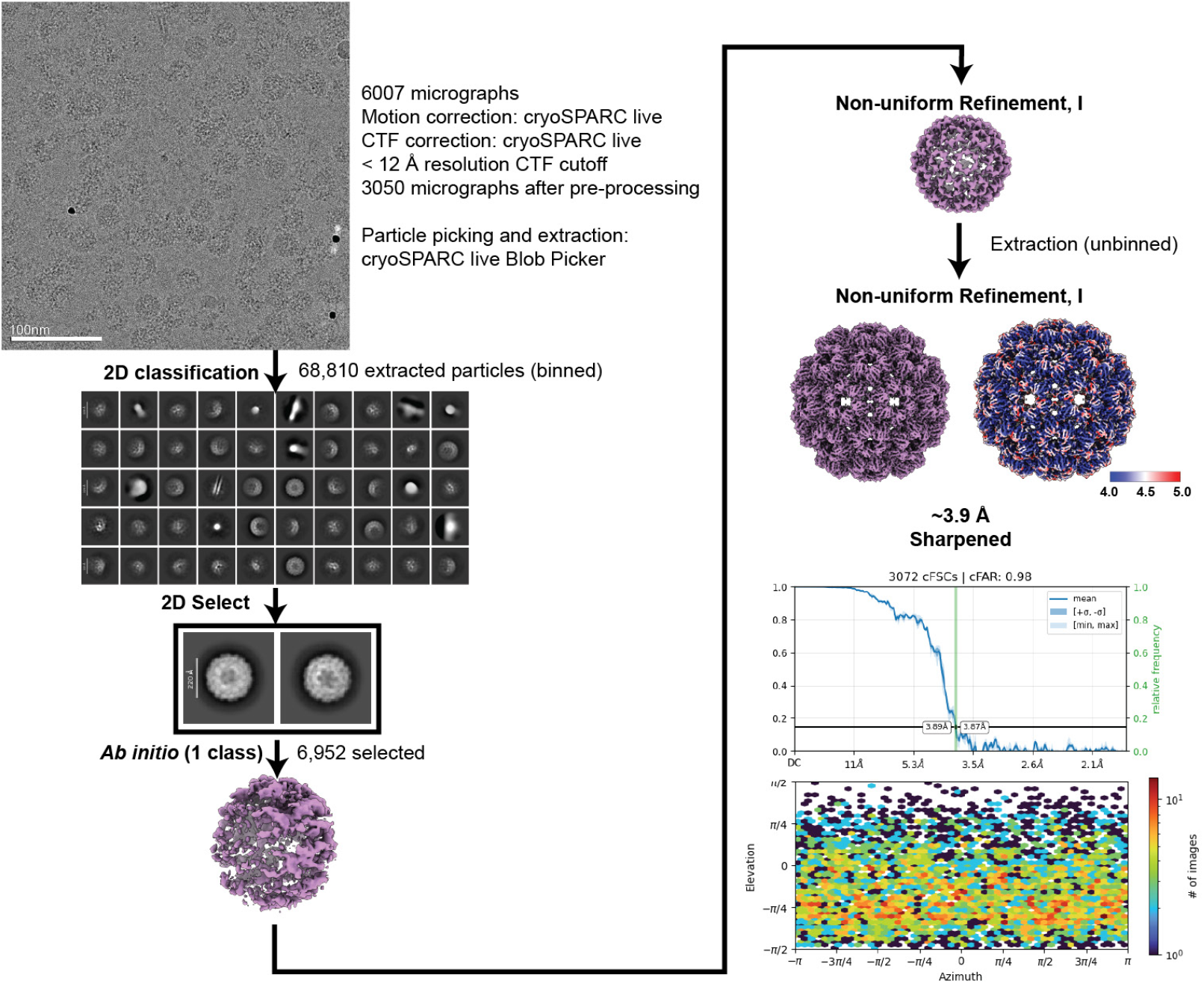
Processing pipeline of the pH-shift CCMV dataset prepared with MIU. Representative processing pipeline for the pH-shift CCMV dataset. Motion and CTF correction were performed in cryoSPARC Live. Particles were picked using blob-based picking, followed by 2D classification and selection of a particle subset for *ab initio* reconstruction. The resulting volume was refined with icosahedral symmetry imposed. After re-extraction, particles were further refined to yield a final map at ~3.9 Å resolution. The final map is shown next to a local resolution map. The conical FSC plot overlaid with a resolution histogram and corresponding angular distribution plot are shown below the final map.

**Figure S9.**
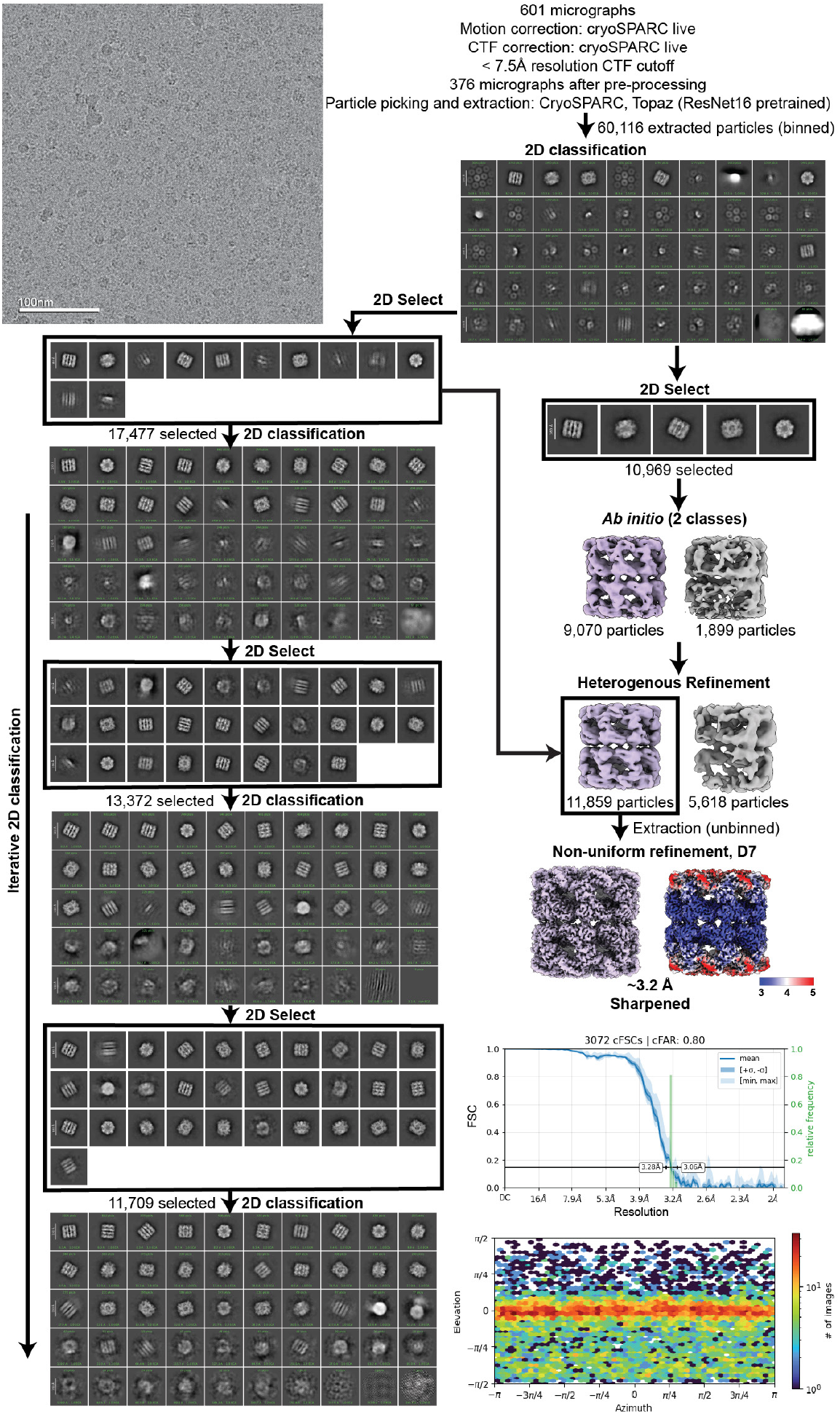
Processing pipeline of the control dataset of GroEL in the presence of GroES and absence of ATP. Representative processing pipeline for the GroEL + GroES control dataset prepared in the absence of ATP. Motion and CTF correction were performed in cryoSPARC Live. Micrographs were exported to cryoSPARC, and particles were picked using Topaz with a pretrained model, followed by initial 2D classification. Iterative 2D classification was also performed to confirm the absence of GroEL/ES complexes. A subset of particles resembling GroEL was selected for *ab initio* reconstruction, and the resulting volumes were further refined in a heterogeneous refinement job using a broader particle set. High-quality particles were then re-extracted and refined with imposed D7 symmetry to yield a final map at ~3.2 Å resolution. The conical FSC plot overlaid with a resolution histogram and corresponding angular distribution plot are shown below the final map.

**Figure S10.**
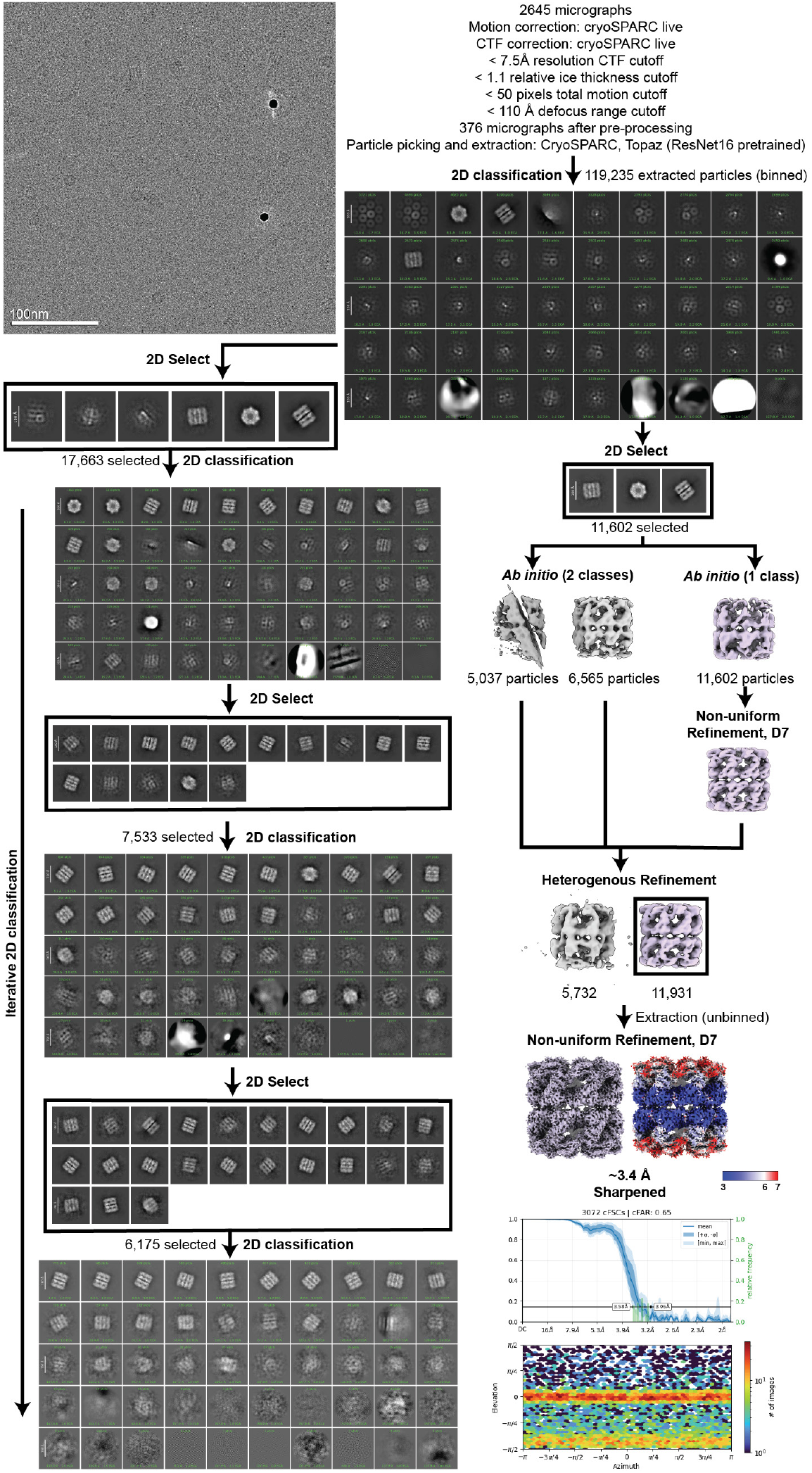
Processing pipeline of the 100 ms GroEL/ES + ATP dataset. Representative processing pipeline for the 100 ms GroEL + GroES + ATP dataset. Motion and CTF correction were performed in cryoSPARC Live. Micrographs were exported to cryoSPARC, and particles were picked using Topaz with a pretrained model, followed by 2D classification and selection of GroEL particles, resembling the “barrel” (uncapped) conformational state. Iterative 2D classification was performed in parallel to confirm the absence of additional conformational states. *Ab initio* reconstructions were generated and the “best” class was further refined with symmetry imposed. Heterogeneous refinement was then used to sort high-quality particles from a broader subset selection. The best particles were re-extracted and refined with D7 symmetry to yield a final map at ~3.4 Å resolution. The final map is shown next to a local resolution map. The conical FSC plot overlaid with a resolution histogram and corresponding angular distribution plot are shown below the final maps.

**Figure S11.**
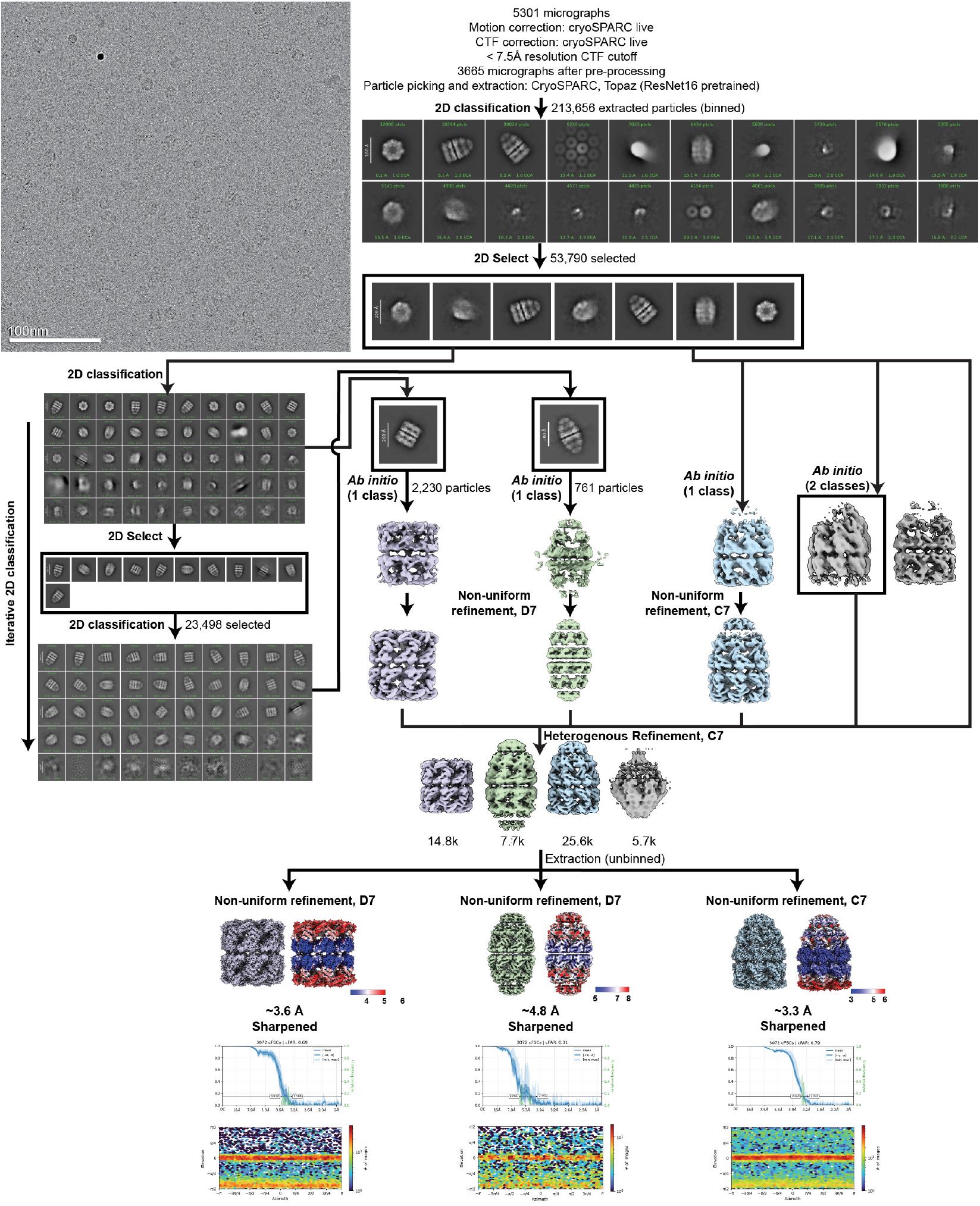
Processing pipeline of the 300 ms GroEL/ES + ATP dataset. Representative processing pipeline for the 300 ms GroEL+ GroES + ATP dataset. Motion and CTF correction were performed in cryoSPARC Live. Micrographs were exported to cryoSPARC, and particles were picked using Topaz with a pretrained model, followed by initial 2D classification and selection of GroEL/ES complexes, all resembling the “bullet” (single-capped) conformation. Further rounds of 2D classification revealed two additional conformational states – the barrel (uncapped), and football (double-capped). *Ab initio* reconstructions were performed independently for each of the three conformational states. These reconstructions, along with a junk volume, were used in heterogeneous refinement with C7 symmetry imposed to further refine particle sets. Sorted particles were re-extracted and refined with symmetry to yield final maps at ~3.3 Å (bullet), ~3.6 Å (barrel), and ~4.8 Å (football) resolution. Final maps are shown next to local resolution maps. The conical FSC plot overlaid with a resolution histogram and corresponding angular distribution plot are shown below the final maps.

**Figure S12.**
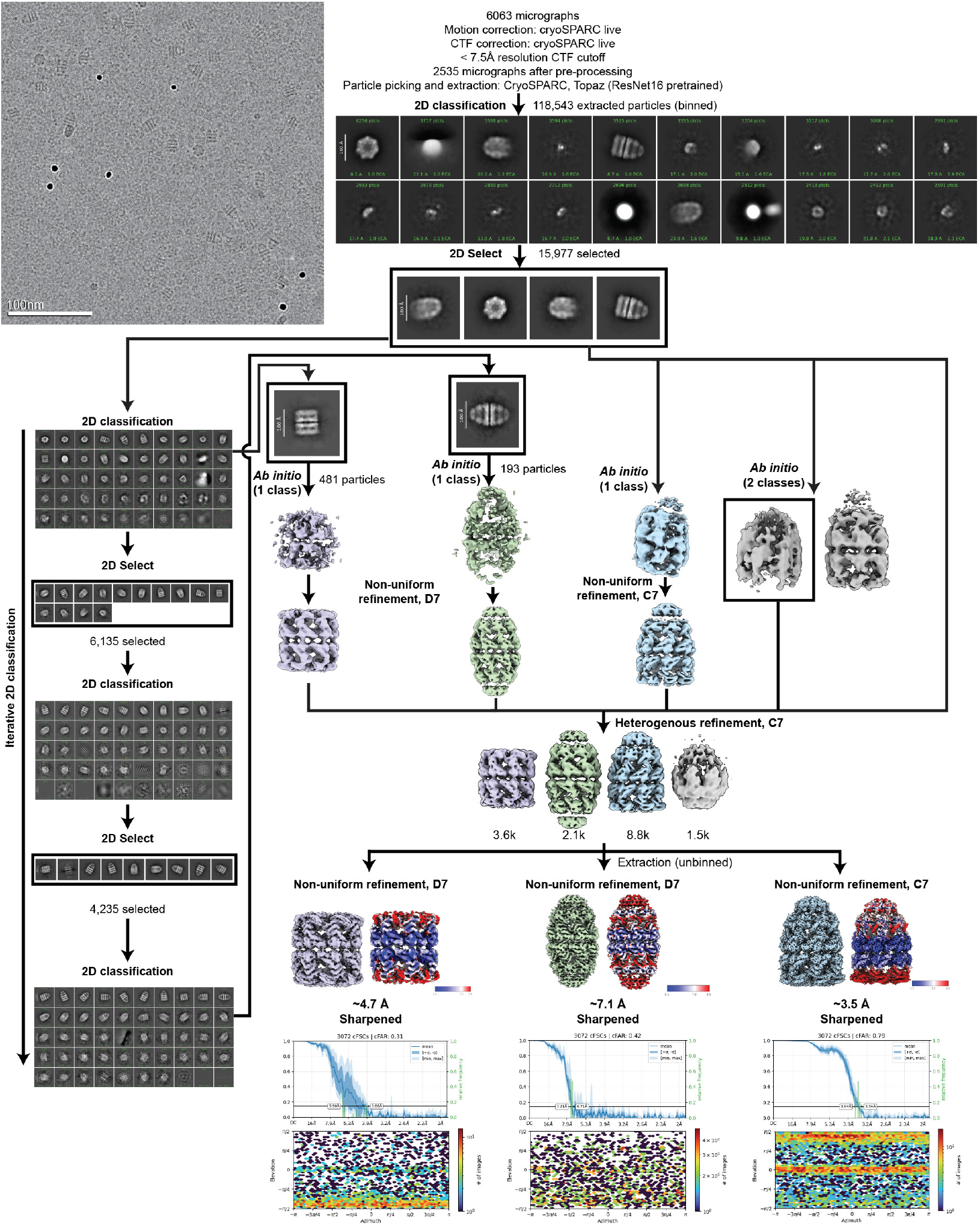
Processing pipeline of the 700 ms GroEL/ES + ATP dataset. Representative processing pipeline for the 700 ms GroEL + GroES + ATP dataset. Motion and CTF correction were performed in cryoSPARC Live. Micrographs were exported to cryoSPARC, and particles were picked using Topaz with a pretrained model, followed by initial 2D classification and selection of GroEL/ES complexes. Further 2D classification revealed three distinct conformational states: bullet (single-capped), barrel (uncapped), and football (double-capped). *Ab initio* reconstructions were performed independently for each conformation, and the resulting volumes were used in a heterogeneous refinement job with C7 symmetry imposed to sort particles. Sorted particles were re-extracted and refined with appropriate symmetry to yield final maps at ~3.5 Å (bullet), ~4.7 Å (barrel), and ~7.1 Å (football) resolution. Final maps are shown next to local resolution maps. The conical FSC plot overlaid with a resolution histogram and corresponding angular distribution plot are shown below the final maps.

**Table S1.**
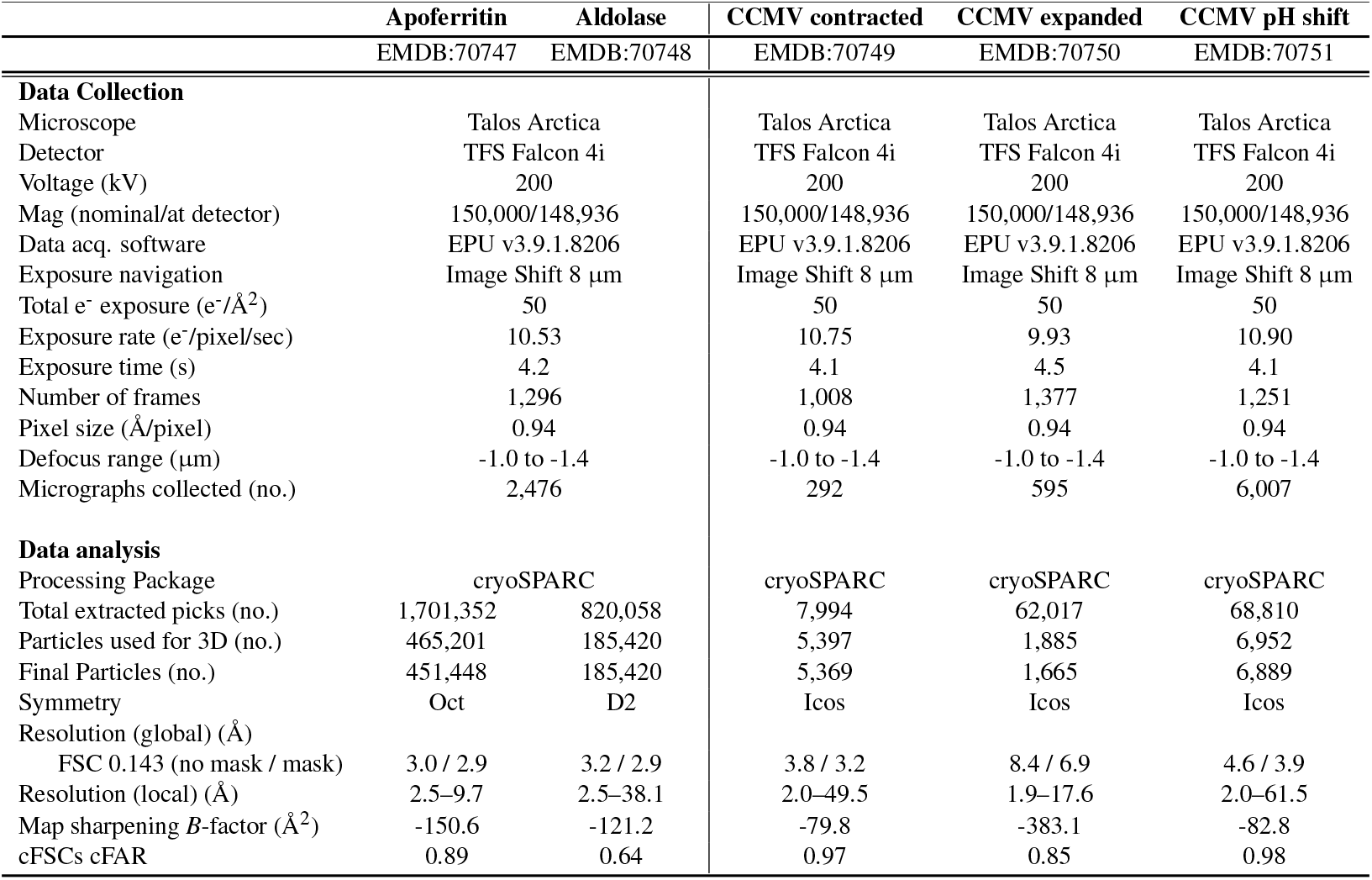
Cryo-EM data collection and image analysis for MIU apoferritin+aldolase, CCMV.

**Table S2.** Cryo-EM data collection and image analysis for MIU GroEL+GroES.

|  | GroEL barrel<br>no ATP | GroEL barrel<br>100 ms | GroEL bullet<br>300 ms | GroEL bullet<br>300 ms | GroEL football<br>300 ms | GroEL barrel<br>700 ms | GroEL bullet<br>700 ms | GroEL football<br>700 ms |
| --- | --- | --- | --- | --- | --- | --- | --- | --- |
|  | EMDB:70752 | EMDB:70759 | EMDB:70753 | EMDB:70754 | EMDB:70755 | EMDB:70756 | EMDB:70757 | EMDB:70758 |
| **Data Collection** |  |  |  |  |  |  |  |  |
| Microscope | Talos Arctica | Talos Arctica |  | Talos Arctica |  |  | Talos Arctica |  |
| Detector | TFS Falcon 4i | TFS Falcon 4i |  | TFS Falcon 4i |  |  | TFS Falcon 4i |  |
| Voltage (kV) | 200 | 200 |  | 200 |  |  | 200 |  |
| Mag (nominal/at detector) | 150,000/148,936 | 150,000/148,936 |  | 150,000/148,936 |  |  | 150,000/148,936 |  |
| Data acq. software | EPU v3.9.1.8206 | EPU v3.9.1.8206 |  | EPU v3.9.1.8206 |  |  | EPU v3.9.1.8206 |  |
| Exposure navigation | Image Shift 8 μm | Image Shift 8 μm | | Image Shift 8 μm | | | Image Shift 8 μm | |
| Total e<sup>-</sup> exposure (e<sup>-</sup>/Å<sup>2</sup>) | 50 | 50 |  | 50 |  |  | 50 |  |
| Exposure rate (e<sup>-</sup>/pixel/sec) | 10.90 | 10.94 |  | 9.66 |  |  | 10.90 |  |
| Exposure time (s) | 4.1 | 4.1 |  | 4.6 |  |  | 4.1 |  |
| Number of frames | 1,251 | 1,251 |  | 1,395 |  |  | 1,251 |  |
| Pixel size (Å/pixel) | 0.94 | 0.94 |  | 0.94 |  |  | 0.94 |  |
| Defocus range (μm) | -1.0 to -1.4 | -1.0 to -1.4 | | -1.0 to -1.4 | | | -1.0 to -1.4 | |
| Micrographs collected (no.) | 601 | 2,645 |  | 5,301 |  |  | 6,063 |  |
| **Data analysis** |  |  |  |  |  |  |  |  |
| Processing Package | cryoSPARC | cryoSPARC |  | cryoSPARC |  |  | cryoSPARC |  |
| Total extracted picks (no.) | 60,116 | 119,235 |  | 255,389 |  |  | 118,543 |  |
| Particles used for 3D (no.) | 11,859 | 11,931 | 14,813 | 25,588 | 7,696 | 3,605 | 8,753 | 2,121 |
| Final Particles (no.) | 11,777 | 11,864 | 14,730 | 25,430 | 7,653 | 3,588 | 8,711 | 2,111 |
| Symmetry | D7 | D7 | D7 | C7 | D7 | D7 | C7 | D7 |
| Resolution (global) (Å) |  |  |  |  |  |  |  |  |
| FSC 0.143 (no mask / mask) | 3.8 / 3.2 | 3.9 / 3.4 | 4.2 / 3.6 | 3.9 / 3.3 | 6.8 / 4.8 | 7.0 / 4.7 | 4.4 / 3.5 | 8.1 / 7.1 |
| Resolution (local) (Å) | 2.0-31.8 | 3.1-41.5 | 4.3-38.2 | 1.9-17.4 | 5.0-51.4 | 2.0-70.5 | 2.0-55.3 | 1.9-17.4 |
| Map sharpening *B*-factor (Å<sup>2</sup>) | -75.4 | -75.2 | -84.7 | -74.3 | -180.9 | -118.4 | -60.5 | -685.9 |
| cFSCs cFAR | 0.80 | 0.65 | 0.69 | 0.79 | 0.31 | 0.31 | 0.79 | 0.42 |

